# A positive feedback loop mediates crosstalk between calcium, cyclic nucleotide and lipid signalling in *Toxoplasma gondii*

**DOI:** 10.1101/2021.11.29.470317

**Authors:** Caia Dominicus, Stephanie D. Nofal, Malgorzata Broncel, Nicholas J. Katris, Helen R. Flynn, Gustavo Arrizabalaga, Cyrille Y. Botté, Brandon M. Invergo, Moritz Treeck

**Affiliations:** Signalling in Apicomplexan Parasites Laboratory, The Francis Crick Institute, 1 Midland Road, NW1 1AT, London, United Kingdom; Protein Analysis and Proteomics Platform, The Francis Crick Institute, 1 Midland Road, NW1 1AT, London, United Kingdom; Apicolipid group, Institut Albert Bonniot UMR5309, CNRS, Université Grenoble Alpes, INSERM, Domaine de la Merci, 38700, La Tronche, France; University of Indianapolis, School of Medicine, 635 Bernhill Drive, 46202 Indianapolis, United States of America; Translational Research Exchange @ Exeter, University of Exeter, Exeter, UK

## Abstract

Fundamental processes of obligate intracellular parasites, such as *Toxoplasma gondii* and *Plasmodium falciparum*, are controlled by a set of plant-like calcium dependent kinases (CDPKs), the conserved cAMP- and cGMP-dependent protein kinases (PKA and PKG), secondary messengers and lipid signalling. While some major components of the signalling networks have been identified, how these are connected remains largely unknown. Here, we compare the phospho-signalling networks during *Toxoplasma* egress from its host cell by artificially raising cGMP or calcium levels to activate PKG or CDPKs, respectively. We show that both these inducers trigger near identical signalling pathways and provide evidence for a positive feedback loop involving CDPK3. We measure phospho- and lipid signalling in parasites treated with the Ca^2+^ ionophore A23187 in a sub-minute timecourse and show CDPK3-dependent regulation of diacylglycerol levels and increased phosphorylation of four phosphodiesterases (PDEs), suggesting their function in the feedback loop. Disruption of CDPK3 leads to elevated cAMP levels and inhibition of PKA signalling rescues the egress defect of ΔCDPK3 parasites treated with A23187. Biochemical analysis of the four PDEs identifies PDE2 as the only cAMP-specific PDE among these candidates, while the other PDEs are cGMP specific, two of which are inhibited by the predicted PDE inhibitor BIPPO. Conditional deletion of the four PDEs supports an important, but non-essential role for PDE1 and PDE2 in growth, with PDE2 controlling A23187-mediated egress. In summary we uncover a positive feedback loop that enhances signalling during egress and links several signalling pathways together.

## Introduction

The Apicomplexa are obligate intracellular parasites that pose a considerable challenge to human and animal health. The most prevalent member of this phylum, *Toxoplasma gondii*, infects virtually all warm-blooded animals, including an estimated 30% of humans worldwide (Pappas, Roussos and Falagas, 2009). To ensure its survival in the host, *Toxoplasma*, like all apicomplexan parasites, must actively invade host cells to initiate replication inside a parasitophorous vacuole. Following several rounds of division, or in response to adverse environmental changes, tachyzoites are triggered to egress from host cells, allowing for subsequent cycles of reinvasion and growth.

At any stage during the replicative cycle, *T. gondii* may be triggered to egress from infected cells in response to deleterious environmental changes. Both extrinsic and intrinsic stimuli play a role in this process. Extrinsic signals include low potassium (K^+^), low pH, and serum albumin (Moudy, Manning and Beckers, 2001; Roiko, Svezhova and Carruthers, 2014; Brown, Lourido and Sibley, 2016), while the accumulation of phosphatidic acid (PA) produced in the parasitophorous vacuole serves as an intrinsic signal to induce natural egress, although this occurs in a more gradual manner after several cycles of endodyogeny (Bisio et al., 2019).

Irrespective of the egress trigger, it is clear that secondary messengers play a key role in driving the process forward once initiated. Indeed, calcium (Ca^2+^) (Carruthers and Sibley, 1999), purine cyclic nucleotides (cGMP and cAMP) (Wiersma et al., 2004; Ono et al., 2008), and phosphatidic acid (PA)(Bullen et al., 2016) have all been implicated (a model is shown in Supp Fig. 1). Upon initiation of egress, migration, or invasion, cytosolic Ca^2+^ levels rise substantially (Lourido and Moreno, 2015). It has been hypothesised that inositol triphosphate (IP3), generated by phosphoinositide phospholipase C (PI-PLC)- mediated cleavage of phosphatidylinositol 4,5-bisphosphate (PIP2), opens an (as yet unidentified) IP3-sensitive channel to release Ca^2+^ from organelles that otherwise sequester Ca^2+^ during immotile replicative stages (Lovett et al., 2002; Garcia et al., 2017). Once released, Ca^2+^ activates a range of effectors, including a group of ‘plant-like’ Ca^2+^- dependent protein kinases (CDPKs) (Billker, Lourido and Sibley, 2009) and proteins involved in vesicle exocytosis (Farrell et al., 2012). PI-PLC-mediated cleavage of PIP2 also leads to the production of diacylglycerol (DAG), which can be interconverted to PA by the apicomplexan-specific DAG-kinase 1 (DGK1). In conjunction with Ca^2+^, PA is believed to play an indispensable role in microneme secretion by interacting with the PA receptor, acylated pleckstrin-homology (PH) domain-containing protein (APH), to facilitate microneme exocytosis (Bullen et al., 2016).

The apicomplexan cGMP-dependent protein kinase (PKG) has been identified as a key regulator of the above-mentioned Ca^2+^ signalling cascade by facilitating the production of IP3 precursors (Brochet et al., 2014; Katris et al., 2020). Moreover, several studies have suggested that PKG, beyond its regulation of the phosphoinositide pathway, may also exert further control over egress by targeting as yet unidentified substrates required for microneme secretion (Brown, Lourido and Sibley, 2016; Brown, Long and Sibley, 2017) In *T. gondii* tachyzoites, the cAMP-dependent protein kinase catalytic subunit 1 (PKAc1), meanwhile, has been proposed to act as a negative regulator of PKG signalling by inhibiting egress induced by parasite-dependent acidification (Jia et al., 2017; Uboldi et al., 2018). PKAc1 has been suggested to indirectly regulate PKG by phosphorylating and possibly activating a putative cGMP phosphodiesterase (PDE) which would result in the degradation of cGMP and a subsequent down-regulation of PKG activity (Jia et al., 2017). It is important to note that the cAMP signalling pathway plays no role in regulating *P. falciparum* egress during the blood stages of infection, and instead appears to be important for mediating invasion (Leykauf et al., 2010; Flueck et al., 2019; Patel et al., 2019).

Although many experimental observations place PKG (Brown, Long and Sibley, 2017; Jia et al., 2017) and phosphoinositide (Bullen et al., 2016) signalling upstream of cytosolic Ca^2+^ flux (and by extension the activation of CDPKs) a further level of interaction between cGMP signalling and CDPK3 has become apparent (Lourido, Tang and David Sibley, 2012). CDPKs, comprised of a serine/threonine kinase domain fused to a calmodulin-like domain, belong to a superfamily of kinases that feature prominently in the Ca^2+^ signalling pathways of plants and some ciliates. Although *Toxoplasma* has numerous CDPK encoding genes (Long, Wang and Sibley, 2016), CDPK1 and CDPK3 have been most extensively studied. CDPK1 has been implicated in microneme exocytosis and the subsequent initiation of gliding motility (Lourido, Tang and David Sibley, 2012), while CDPK3 has been shown to be important for rapid Ca^2+^ionophore-induced egress, where the addition of the calcium ionophore A23187 or BIPPO leads to concerted parasite exit from the host cell in seconds (Black and Boothroyd, 2000; Garrison et al., 2012; Lourido, Tang and David Sibley, 2012; McCoy et al., 2012). Intriguingly, while a marked delay in egress is evident when CDPK3 depleted/inhibited parasites are treated with Ca^2+^ ionophore, this phenotype is partially rescued when tachyzoites are induced to egress with PDE inhibitors such as zaprinast and BIPPO (Lourido, Tang and David Sibley, 2012; Howard et al., 2015). The Ca^2+^ ionophore A23187 forms lipid-soluble complexes with divalent cations and is thought to induce a discharge of Ca^2+^ from cytosolic stores. Both zaprinast and BIPPO, known inducers of *Toxoplasma* egress (Lourido, Tang and David Sibley, 2012; Howard et al., 2015) are thought to induce an elevation in cytosolic cGMP levels by inhibiting cGMP hydrolysing PDEs, activating PKG activity via elevated cGMP levels. Accordingly, these findings suggest a compensatory role for PKG signalling in the absence of CDPK3.

While this compensatory mechanism has not been examined in any great detail, it is possible that PKG and CDPK3 substrate specificity may overlap. Multiple kinases converging on shared targets can provide multiple layers of regulation to a single pathway, and this is a known feature of nucleotide-activated kinases, including PKG (Pearce, Komander and Alessi, 2010). Alternatively, it is plausible that BIPPO’s compensatory effects are explained by a more direct link between CDPK3 and PKG activity; if CDPK3 were to play a feedback-mediated role in the positive regulation of PKG signalling, pharmacological activation of PKG (e.g. by BIPPO/zaprinast treatment) would also diminish the requirement for CDPK3 during egress. Interestingly, this is reminiscent of the *P. falciparum* CDPK5, where the egress block of CDPK5-deficient parasites can be rescued by hyperactivation of PKG (Absalon et al., 2018).

While the above literature forms a common understanding that CDPKs, PKG, PKA, lipid and second messenger signalling are important across lifecycle stages in *Toxoplasma* and *Plasmodium* species, how they are spatially and temporally regulated, how they intersect and how specific signalling outcomes are achieved is not well described.

Here, we report on the phospho-, lipid and cyclic nucleotide signalling networks activated during the pharmacological induction of *Toxoplasma* tachyzoite egress using either A23187 or BIPPO. Collectively, our data highlights the presence of a feedback loop between A23187-regulated Ca^2+^ release and cyclic nucleotide as well as phosphoinositide signalling. This mechanism appears to be regulated, at least in part, by CDPK3 and the cAMP-specific phosphodiesterase PDE2.

## Results

### Generation of Calcium Reporter Lines to align BIPPO and A23187 signalling pathways

To investigate how the cGMP and calcium signalling pathways converge and differ, we compared their phosphorylation dynamics using two activators of these pathways: BIPPO, a PDE inhibitor, and the calcium ionophore A23187.

The signalling kinetics following Ca^2+^ ionophore and BIPPO treatment vary, so we first determined a timepoint at which both pathways should be comparable. Common to both treatments is a raise in intracellular calcium levels before egress. We therefore chose peak intracellular calcium levels as a reference point to facilitate a direct comparison between BIPPO- and A23187-treated parasites. To this end, we generated a stable calcium sensor line that co-expresses, through use of a T2A ribosomal skip peptide, an internal GFP control and the genetically encoded ruby Ca^2+^ biosensor jRCaMP1b (Alves et al., 2021) from a single promoter (Fig. 1A-B). The expression of the biosensor did not have any discernible effects on Ca^2+^ ionophore (A23187) or BIPPO induced egress rates (Supp Fig. 2) therefore all subsequent experiments were performed with this line (henceforth referred to as WT). While some variability of jRCaMP1b fluorescence was observed between vacuoles at a per-well level upon stimulation, ratiometric quantitation of jRCaMP1b fluorescence upon BIPPO or A23187 treatment of cytochalasin D- immobilised parasites illustrated distinct Ca^2+^ response curves; BIPPO treatment led to a rapid increase in Ca^2+^ levels, (Fig. 1Ci), while the cytosolic Ca^2+^ rise detected upon A23187 treatment appeared more gradual (Fig. 1Cii). To facilitate optimal alignment, and to account for the rapid kinetics of these signalling pathways, treatment timings of 15s (BIPPO) and 50s (A23187) were selected for subsequent phosphoproteomics experiments.

**Figure 1.**
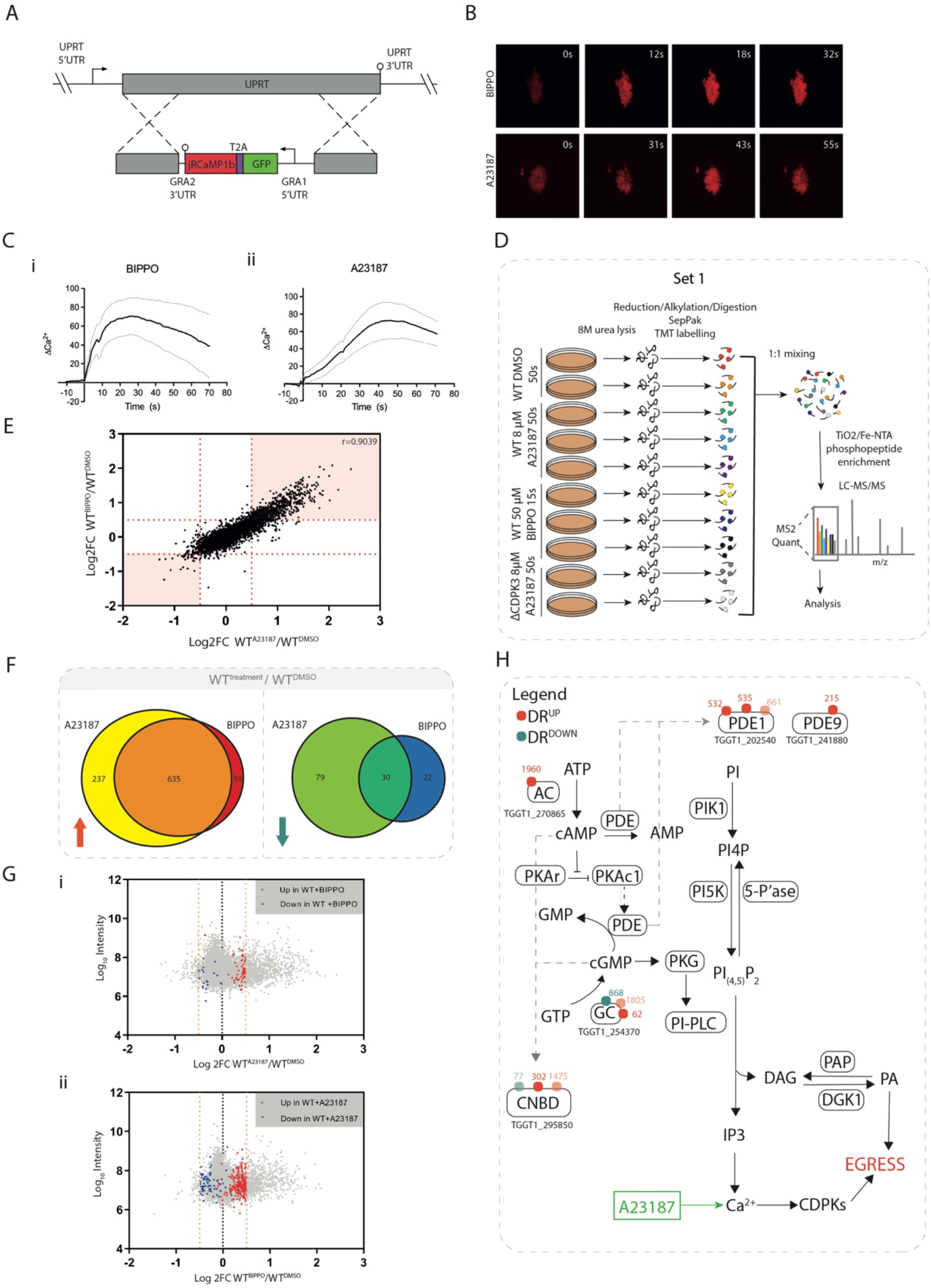
Comparative phosphoproteomics of BIPPO vs A23187-induced GFP-T2A-jRCaMP1b (WT) parasites at peak cytosolic calcium levels (A) Generation of the calcium sensor line GFP-T2A- jRCaMP1b by integration into the UPRT locus. (B) Video microscopy of intracellular GFP-T2A-jRCaMP1b parasites (red channel), following addition of 50 µM BIPPO or 8 µM A23187 at 0s. (C) Ratiometric tracking of mean Ca^2+^ response (jRCaMP1b/GFP normali<u>s</u>ed to 0), following addition of (i) 50µM BIPPO or (ii) 8 µM A23187. Grey dotted lines represent ± SEM. Red dotted line indicates the timepoint selected for subsequent phosphoproteomic experiments. Data was collected from ≥ 10 vacuoles (in separate wells) over ≥6 days. (D) Schematic summary of the TMT-10-plex experiment (set 1) and workflow used to quantify the phosphoproteomes of intracellular WT tachyzoites treated with 50 µM BIPPO (15s) or 8 µM A23187 (50s). ΔCDPK3 parasites treated with 8 µM A23187 (50s) were included to facilitate later analyses. (E) Correlation of phosphosite regulation (log2FC) in BIPPO (y axis) and A23187 (x axis)-treated WT parasites. Each data point corresponds to a single phosphosite. Red dotted lines represent 3xMAD outlier thresholds used to determine differential site regulation. Phosphosites that fall within the pink shaded regions are differentially regulated upon treatment with both BIPPO and A23187. (F) Overlap between phosphosites that are differentially regulated in GFP-T2A-jRCaMP1b (WT) tachyzoites following treatment with 8 µM A23187 (50s) or 50 µM BIPPO (15s). Red arrow represents up-regulated sites (log2FC>0.5), blue arrow represents down-regulated sites (log2FC<-0.5). Only sites confidently detected in both treatment conditions were included for analysis. (G) Differential phosphosite regulation (log2FC) vs log10 total reporter intensity following treatment of WT tachyzoites with (i) 8 µM A23187 (50s) or (ii) 50 µM BIPPO (15s). (H) Each data point corresponds to a single phosphosite. Only sites confidently detected in both BIPPO and A23187 samples are shown. In (i) Coloured data points highlight sites that were differentially up- or down-regulated (red and blue, respectively) upon BIPPO, but not A23187 treatment, while in (ii) coloured data points highlight sites that were differentially up- or down-regulated (red and blue, respectively) upon A23187, but not BIPPO treatment. Orange dotted lines represent 3xMAD outlier thresholds used to determine differential site regulation (log2FC>0.5 for up-regulated sites and log2FC<-0.5 for down-regulated sites). (H) Differentially A23187-regulated phosphosites detected on targets implicated in the regulation of cyclic nucleotides. Red and blue dots represent sites that are differentially up- or down-regulated, respectively. Numbers refer to site position within protein. Dots with reduced opacity represent sites with a localization score difference<10.

### A23187- and BIPPO-treated Wildtype Parasites Exhibit Highly Correlative Phosphoproteomic Responses at temporally aligned calcium flux

Having identified the optimal BIPPO and A23187 treatment timings to achieve maximal calcium release, we wanted to identify and compare phosphorylation events that take place during BIPPO- and A23187-induced signalling cascades at these timepoints. We used multiplexed tandem-mass-tags (TMT) and LC-MS/MS to quantify the phosphoproteome of intracellular WT tachyzoites treated with vehicle (DMSO), 50 µM BIPPO (15s) or 8 µM A23187 (50s) at 37°C. DMSO-treated parasite samples were generated in 2 biological replicates, while BIPPO- and A23187-treated samples were generated in 3 biological replicates (Fig. 1D). At these timepoints parasites remained intracellular. Samples were lysed, digested, and labelled with different TMT tags. Labelled samples were pooled and subjected to TiO2/Fe-NTA phosphopeptide enrichment prior to LC-MS/MS analysis. Of note, this experiment (Set 1), also contains two biological replicates of an A23187-treated calcium-dependent kinase 3 deletion (ΔCDPK3) parasite line. This allowed us to identify ΔCDPK3 dependency of signalling events during A23187 and BIPPO induced egress and is explained further below.

We quantified changes in phosphorylation states by calculating the log2-transformed intensity ratios (log2FC) of A23187- or BIPPO-stimulated WT parasites versus DMSO-treated WT parasites (DataS1). In total we quantified 7,811 phosphorylation sites across these conditions.

Differentially regulated (DR) phosphorylation sites were selected if the log2FC exceeded 3x the median absolute deviation (MAD), rounded to the nearest tenth. This was log2FC>0.5 for up-regulated sites and log2FC<-0.5 for down-regulated sites; (Supp Fig. 3A) and applied across all datasets.

The rapid signalling progression upon treatment with BIPPO and A23187 inevitably results in variability in phosphosite intensities between replicates, where despite our best efforts, signalling may be stopped with several seconds difference between experiments. As such variability results in poor p-values in classical t-tests and, by extension, an under-reporting of true treatment-regulated sites, we did not subject DR sites to further p-value<u>-</u> based thresholding. However, the reporter intensities associated with DR sites correlated well across replicates (r>0.89, Supp Fig. 3B). This suggests that despite some of the aforementioned replicate variability, the overall trends across replicates were consistent, and these scores could therefore be confidently averaged to provide values that are representative of a site’s phosphorylation state at the timepoint of interest.

Comparison of the log2FCs observed in BIPPO- and A23187-treated samples shows strong correlation between the phosphorylation responses of these conditions (r=0.9039) (Fig. 1E), suggesting that the signalling pathways at these selected timepoints align sufficiently well to directly compare them.

### Phosphoproteomic Analysis Cannot Confidently Distinguish BIPPO- from A23187- Induced Signalling

To investigate the signalling events that are shared between or are unique to BIPPO and A23187 treatment, we identified DR sites for each treatment condition. We then identified DR sites that were successfully quantified in both treatments, which allowed us to examine their behaviour under both conditions. In total we identified 746 BIPPO and 981 A23187 DR sites. A large overlap was detected between treatments for both up- and down-regulated phosphosites (DR^UP^ and DR^DOWN^, respectively); ∼91% of phosphosites up-regulated following BIPPO treatment showed similar regulation upon A23187 addition and ∼58% of BIPPO down-regulated sites behaved similarly following A23187 treatment (Fig. 1F).

We also observed some dissimilar regulation between conditions; 59 phosphorylation sites were found to be up-regulated following BIPPO treatment only, while 237 sites were phosphorylated exclusively following A23187 treatment. Of the DR^DOWN^ phosphorylation sites, 22 were found to be unique to BIPPO treatment, while 79 were unique to A23187 treatment. These treatment-specific sites may originate from distinct signalling pathways, activated by each of the compounds. To discern whether these disparate site behaviours are truly treatment-specific effects, or whether they are the result of imperfect alignment of the treatment timings, we visuali<u>s</u>ed phosphorylation site log2FCs following A23187 treatment, and highlighted phosphorylation sites that were only DR following BIPPO treatment (Fig. 1Gi). Similarly, we also visualised phosphorylation site log2FCs following BIPPO treatment, and highlighted phosphorylation sites that were only DR following A23187 treatment (Fig. 1Gii). In both instances, most sites approached the DR thresholds for up- or down-regulation. While this does not preclude the possibility that some of the BIPPO- and A23187-specific DR^UP/DOWN^ sites are regulated in a drug-exclusive manner, it is likely that the majority of these sites would pass the DR threshold within seconds, and that minor changes in treatment timing can make the difference between surpassing the DR threshold or not.

Collectively, these findings demonstrate that at temporally aligned calcium release within the parasite, it is nearly impossible to detect clear signalling features that confidently distinguish the BIPPO-activated signalling pathway from the signalling cascade activated upon cytosolic Ca^2+^ elevation by A23187 treatment.

### A23187 Treatment Leads to Differential Regulation of Targets Implicated in the PKG Signalling Pathway

A substantial overlap between BIPPO-regulated and A23187-regulated sites was expected given the increase of cytosolic Ca^2+^ in both treatment conditions. However, the inability to confidently distinguish BIPPO from A23187 signalling was unexpected, as these agents are believed to initiate egress by activating distinct, albeit interconnected, signalling responses (Lourido, Tang and David Sibley, 2012). Previous studies have placed PKG activation upstream of Ca^2+^ release (Brochet et al., 2014; Stewart et al., 2017) and it was therefore to our surprise that within the A23187 and BIPPO response overlap, DR phosphorylation sites were detected on proteins implicated in the catalysis and hydrolysis of the cyclic nucleotides (cNMPs) cGMP and cAMP, key molecules involved in PKG activation. Subsequent examination of all A23187-regulated phosphorylation sites (including those for which we lacked quantifications in BIPPO samples) identified differential phosphorylation on proteins including, but not limited to, enzymes important for cNMP signalling: PDEs (TGGT1_202540 (PDE1) and TGGT1_241880 (PDE9)), an adenylate cyclase (TGGT1_270865), a guanylyl cyclase (TGGT1_254370), and a cyclic nucleotide (cNMP) binding domain (CNBD) containing protein (TGGT1_295850) (Fig. 1H).

The differential phosphorylation of several proteins in the upstream pathway of cNMP production/regulation hints at a putative Ca^2+^-mediated feedback loop that regulates cGMP and/or cAMP signalling. The existence of such a feedback mechanism could account for our inability to confidently discern PKG-specific signalling upon BIPPO treatment, as such signalling would be activated upon treatment with both BIPPO and A23187.

### Deletion of CDPK3 leads to signalling perturbations in both A23817 and BIPPO treatment conditions

Following our analyses of BIPPO- and A23187- induced egress, we set out to explore the role of CDPK3 in these signaling pathways. The observation that BIPPO/zaprinast- mediated activation of PKG partially compensates for a loss of CDPK3 has led to the hypothesis that, given the function of both kinases in egress, the kinases’ substrate specificities may overlap (Lourido, Tang and David Sibley, 2012). In such a scenario, BIPPO treatment would facilitate PKG-mediated phosphorylation of CDPK3 targets, thus overcoming the egress delay otherwise seen in A23187-treated parasites.

To identify phosphorylation sites that might fit such criteria, we wanted to identify phosphorylation sites that are CDPK3-dependent upon A23187s treatment, but CDPK3- independent upon BIPPO treatment. To do this, we generated a ΔCDPK3 parasite line by replacing the endogenous CDPK3 locus in the RH GFP_T2A_jRCaMP1b line with a HXGPRT expression cassette (henceforth known as ΔCDPK3; Fig. 2A) and confirmed deletion of CDPK3 by PCR (Supp Fig. 4). We performed an egress assay to validate the known A23187-induced egress delay reported for ΔCDPK3 parasites (Black and Boothroyd, 2000; Garrison et al., 2012; Lourido, Tang and David Sibley, 2012; McCoy et al., 2012). As expected, we found that A23187-induced egress was substantially inhibited in this line (Fig. 2Bi), while a less severe egress delay was observed in BIPPO-treated ΔCDPK3 parasites (Fig. 2Bii). This recapitulates previous findings (Lourido, Tang and David Sibley, 2012) that activation of PKG partially compensates for a loss of CDPK3.

**Figure 2.**
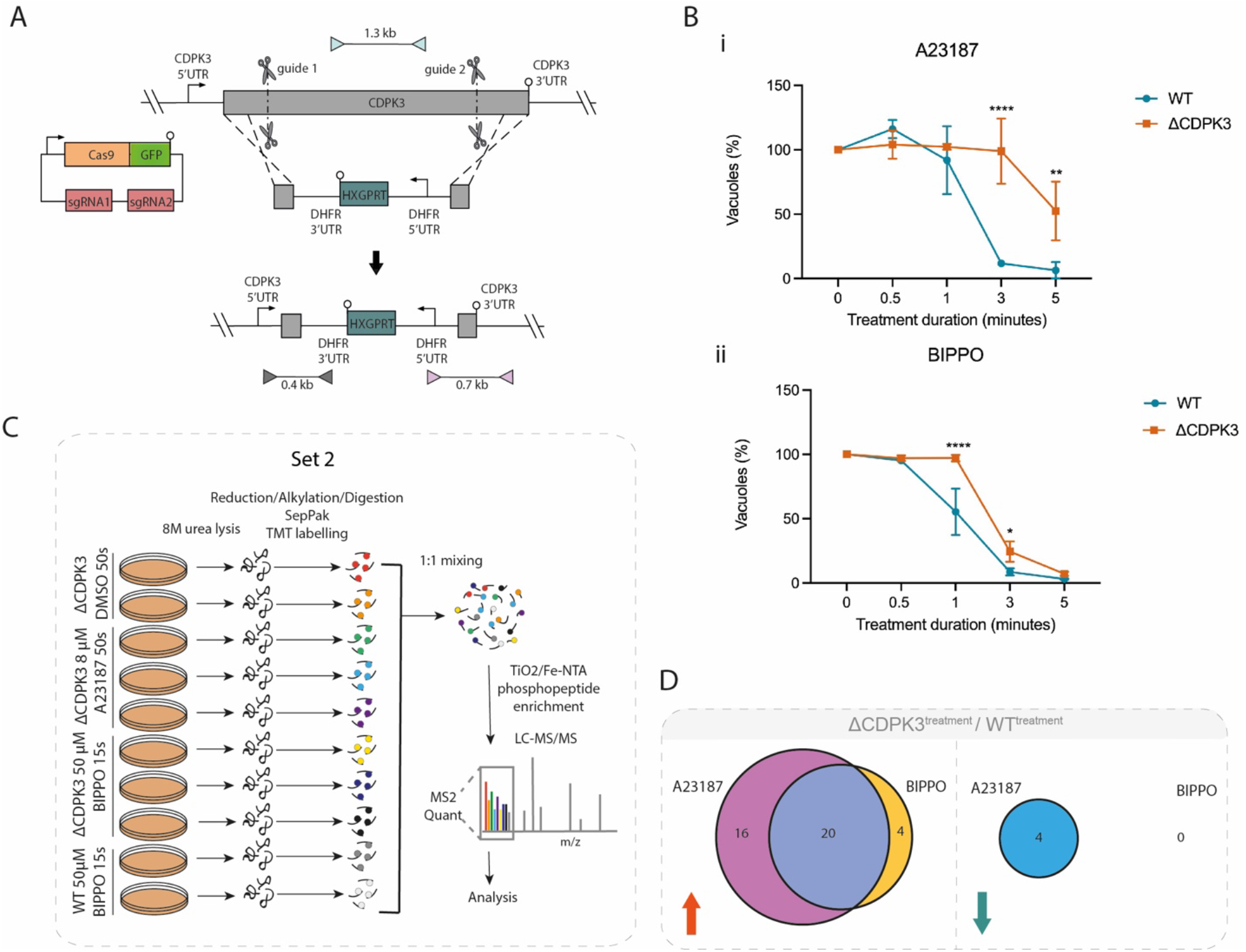
Comparative phosphoproteomics of BIPPO vs A23187-induced GFP-T2A-jRCaMP1b ΔCDPK3 parasites at peak cytosolic calcium levels. **(A)** Generation of the GFP-T2A-jRCaMP1b ΔCDPK3 line using CRISPR/Cas9 to increase site-directed integration. Scissors represent Cas9 cleavage sites and lollipops depict stop codons. Coloured triangles represent primer pairs used to detect WT, 5’ integration and 3’ integration loci (light blue, grey and pink respe<u>c</u>tively). PCR results using these primer pairs are shown in Supp Fig. 4 **(B)** Egress assay of GFP-T2A-jRCaMP1b (WT) and GFP-T2A-jRCaMP1b ΔCDPK3 (ΔCDPK3) parasites following treatment with **(i)** 8 µM A23187 or **(ii)** 50µM BIPPO. Graph shows the remaining % of un-egressed vacuoles (relative to untreated) following A23187/BIPPO treatment. Data are represented as mean ± s.d. (n=3). Two-way ANOVA with Holm-Sidak *post hoc* comparison.****, *P* ≤ 0.0001 **(C)** Schematic summary of the TMT-10-plex experiment (set 2) and workflow used to quantify the phosphoproteomes of intracellular ΔCDPK3 tachyzoites treated with 50 µM BIPPO (15s), 8 µM A23187 (50s), and WT parasites treated with 50 µM BIPPO (15s). **(D)** Overlap between differentially regulated phosphosites that display CDPK3 dependency following treatment with 50 µM BIPPO (15s) or 8 µM A23187 (50s) (data derived from set 1 and 2). Red and blue arrows represent up- and down-regulated sites, respectively. Only phosphosites found to be differentially regulated upon treatment with both A23187 and BIPPO were included for analysis.

We next quantified phosphorylation events in ΔCDPK3 parasites treated with DMSO, 50 µM BIPPO (15s) or 8 µM A23187 (50s) at 37°C (Fig. 2C) in biological replicates (2x DMSO, 3x A23187, 3x BIPPO). In this experiment (set 2), we included 2 biological replicates of BIPPO-treated WT parasites. In conjunction with the ionophore-treated ΔCDPK3 parasites included in set 1 (Fig. 1D), this allowed us to identify CDPK3- dependent phosphorylation sites during BIPPO- and A23187-induced egress (Data S2). We first identified DR phosphorylation sites across all datasets for which we had quantifications and tested for CDPK3 dependency. Of the 498 phosphosites detected, 44 sites (∼8.5%) were CDPK3-dependent (Fig. 2D, Data S2). 40 sites were classed as DR^UP^; 16 were exclusive to A23187 treatment, 20 were identified upon both A23187 and BIPPO treatment, and 4 were detected upon BIPPO treatment only. By contrast, only 4 sites were classed as DR^DOWN^, all in a seemingly A23187-exclusive manner.

The 16 phosphorylation sites that show CDPK3 dependency exclusively upon A23187 treatment constituted putative candidates for PKG/CDPK3 substrate overlap. We reasoned that, if a DR^UP^ site was found to be CDPK3-dependent upon A23187 treatment only, this phosphorylation site should be recovered in ΔCDPK3 parasites following BIPPO treatment. Only 3 of these phosphorylation sites showed this behaviour and were located on two hypothetical proteins (TGGT1_243460, TGGT1_232120) and a DnaJ domain-containing protein (TGGT1_203380) (Data S2). While these findings do not completely rule out the ‘substrate overlap’ theory to account for BIPPO’s compensatory effects, the putative overlap is extremely small, and none of these proteins contain predicted domains that would explain the rescue of CDPK3 mutants by BIPPO in induced egress.

Collectively, our current findings provide more evidence for a Ca^2+^-regulated feedback loop model than for PKG/CDPK3 substrate overlap.

### A sub-minute timecourse of CDPK3 dependent and independent signalling progression in ionophore-induced egress

The experiments delineated above, exploring the phosphosignalling networks in WT and ΔCDPK3 parasites triggered upon treatment with A23187 or BIPPO, examined only a single treatment timepoint. These experiments therefore offer only a limited ‘signalling snapshot’ that precludes any insights into the dynamics of signalling progression over time. To further investigate the putative activation of cNMP-induced signalling pathways in WT and ΔCDPK3 mutants upon A23187 treatment we performed a sub-minute phosphosignalling timecourse. We treated intracellular WT and ΔCDPK3 tachyzoites with 8 µM A23187 for 15, 30 or 60 seconds at 37°C (Fig. 3A), during which the parasites remained intracellular. As before, samples were subjected to TMT-based quantitative analysis of phosphoproteomic changes. Fold changes were calculated relative to a 0s (DMSO) control. In total we quantified 11,021 phosphorylation sites (Data S3).

**Figure 3.**
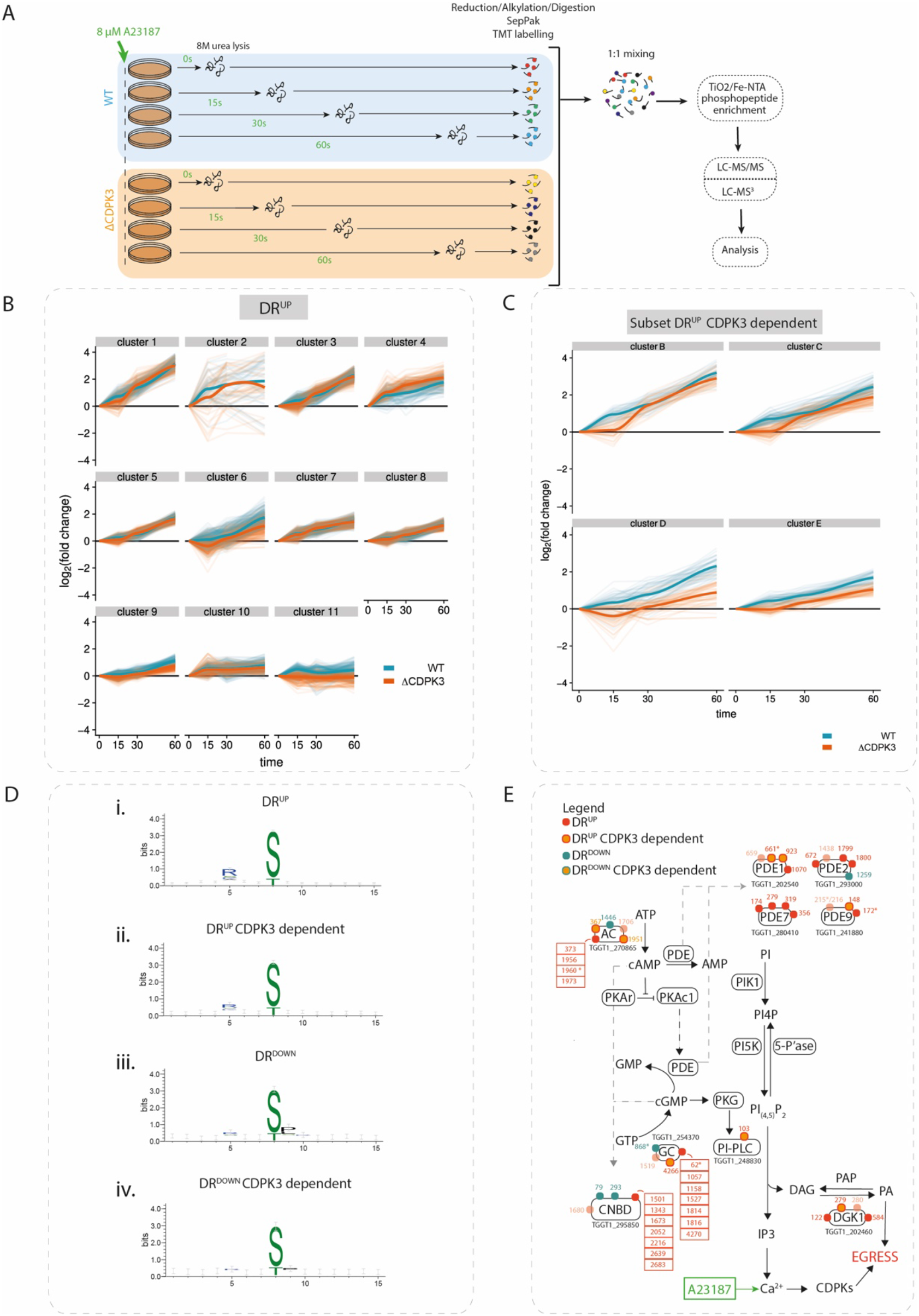
A23187 treatment results in partially CDPK3-dependent phosphorylation of targets implicated in PKG signalling. **(A)** Schematic of A23187-treatment timecourse experimental design. **(B)** Gaussian mixture-model-based clustering of all DR^UP^ sites in the A23187-treatment time courses. Log2FC values from both WT and ΔCDPK3 samples were combined to cluster on six dimensions (WT 15s, 30s and 60s and ΔCDPK3 15s, 30s and 60s). Thin lines represent the time course traces of individual phosphorylation sites. Thick lines represent Loess regression fits of all traces. **(C)** Gaussian mixture-model-based clustering of a subset of DR^UP^ CDPK3-dependent sites in the A23187-treatment time courses. Log2FC values from both WT and ΔCDPK3 samples were combined to cluster on six dimensions (WT 15s, 30s and 60s and ΔCDPK3 15s, 30s and 60s). Thin lines represent the time course traces of individual phosphorylation sites. Thick lines represent Loess regression fits of all traces. Four clusters (B-E) best illustrating the transient phosphorylation delay in ΔCDPK3 parasites are shown. **(D)** Results of phosphorylation motif enrichment analysis using rmotifx 1.0. Results are shown for the analysis of **(i)** DR^UP^ **(ii)** DR^UP^ CDPK3-dependent **(iii)**DR^DOWN^ and **(iv)** DR^DOWN^ CDPK3-dependent phosphorylation sites. **(E)** Differentially A23187-regulated phosphosites detected on targets implicated in the regulation of PKG signalling. Red and blue dots represent sites that are differentially up- or down-regulated, respectively. Dots with an orange centre indicate CDPK3-dependent sites. Numbers refer to site position within protein. Dots with reduced opacity represent sites with a localisation score difference<10. Asterisks highlight sites that were previously detected in this study’s A23187/BIPPO phosphoproteome experiment (see Fig. 1D& Fig. 2C).

DR thresholds were set at 3x MAD of the log2FC across each WT timepoint (15s, 30s and 60s). Phosphorylation sites were considered differentially regulated if at any given timepoint their log2FC surpassed these thresholds. CDPK3 dependency was determined for each phosphorylation site by calculating the log2 ratios of A23187-treated WT and ΔCDPK3 parasites for each timepoint. The resulting ratios were used to calculate the MAD at each timepoint, and the most stringent score was used to set 3X MAD outlier thresholds. A DR site was considered CDPK3-dependent if, at any given timepoint, it simultaneously passed the DR and CDPK3 dependency thresholds.

We identified 2,408 phosphorylation sites (DR^UP^) upon A23187 treatment in WT parasites, which were also quantified in ΔCDPK3 parasites (Data S3). To examine whether this dataset recapitulates our previous findings, we compared the DR^UP^ sites identified at the 60s timepoint in this experiment, with those identified after 50s A23187 treatment from the preceding experiments (Fig. 2C). Of the 572 DR phosphorylation sites identified 50s after A23187 treatment in our initial experiment, 503 sites (∼88%) also passed the threshold for differential (up)regulation in the timecourse at the 60s post-treatment timepoint. Reassuringly, we observed many proteins previously identified as being phosphorylated in a CDPK3-dependent manner (Treeck et al., 2014; Wallbank et al., 2019), including the CRAL/TRIO domain containing protein (TGGT1_254390), a putative P-type ATPase4 (TGGT1_278660), CDPK2A (TGGT1_206590) and the tyrosine transporter ApiAT5-3 (TGGT1_257530).

To get an overview of the progression of signalling cascades in DMSO- and A23187- treated parasites, we performed a clustering analysis, as previously described (Invergo et al., 2017), of DR phosphorylation sites identified in WT and, separately, of DR sites found to be CDPK3-dependent in our timecourse experiments (see Table S1 for all clusters). We obtained 11 clusters showing distinct up-regulation dynamics (Fig. 3B) and 10 clusters showing down-regulation dynamics in WT parasites (Supp Fig. 5A). Analysis of CDPK3-dependent DR sites, meanwhile, yielded 10 up-regulated clusters and 6 down- regulated clusters Fig. 3C), (Supp Fig. 5B-C).

In the up-regulated clusters, we identified a preponderance for phosphorylation motifs with arginine in the -3 position (Fig. 3Di), a consensus sequence that has previously been shown to be preferentially phosphorylated by CDPK1 (Lourido et al., 2013) and possibly CDPK3 (Treeck et al., 2014). Reassuringly, this consensus motif was also observed among DR^UP^ CDPK3-dependent sites (Fig. 3Dii). Down-regulated phosphorylated sites, meanwhile, show a clear enrichment for proline in the +1 position (Fig. 3Diii & Fig. 3Div). This indicates that while CDPK activity (and/or activity of kinases with a similar substrate preference) is being induced by calcium-signalling, a distinct set of one or more kinases with this phosphorylation motif preference is being inactivated concurrently. Alternatively, this could be mediated by the activation of a specific phosphatase. We observed that several of the less phosphorylated proteins in the timecourse are secreted into the parasitophorous vacuole (PV), which physically separates the parasite from the host cell cytoplasm. Several proteins that are secreted into the PV have been shown to play a role in mediating egress. This includes GRA41, which has been shown to be important for A23187-induced egress (LaFavers et al., 2017). It is therefore possible that the secreted proteins identified in our timecourse may be implicated in wider signalling events occurring in the PV that are required for egress.

Several functionally related proteins were phosphorylated with similar dynamics, as revealed by Gene Ontology (GO) term enrichment (see Tables S2-5 for DR^UP^, DR^DOWN^, CDPK3-dependent DR^UP^ and CDPK3-dependent DR^DOWN^ GO term enrichments, respectively). Most notably, two up-regulation clusters (clusters 1 and 2) were enriched in terms related to signal transduction (GO:0007165, GO:0007154, GO:0023052) and hydrolase activity (GO:0016787, GO:0042578, GO:0016462, GO:0016817, GO:0016818, GO:0017111), respectively (Fig. 3B). These enrichments were, in part, driven by phosphorylation of PDEs and cyclases involved in cyclic nucleotide signalling. Thus, not only are the enzymes potentially upstream of PKG being phosphorylated upon exposure to ionophore, but also the dynamics of phosphorylation are similar between them.

We also found significant enrichment of membrane proteins in CDPK3-dependent clusters (GO:0044425, GO:0016021, GO:0031224, GO:0016020). These are predicted to play roles in nutrient transport and ion-exchange, including the sodium-hydrogen exchangers NHE1 and NHE3 (which have previously been linked to egress) (Arrizabalaga et al., 2004; Francia et al., 2005), and the tyrosine transporter ApiAT-5-3 (a known target of CDPK3 phosphorylation)(Treeck et al., 2014; Wallbank et al., 2019).

Further GO enrichment analysis of up-regulated clusters revealed other potential downstream targets of ionophore-induced signalling, including transcription (e.g. GO:0006355, GO:1903506, GO:2001141) by AP2-family transcription factors, magnesium chelatase activity (GO:0016851) by DNA replication licensing factors, and regulation of a GTPase-mediated process (GO:00423087, GO:0051336). Intriguingly, we found that phosphorylation of several targets associated with these GO terms also showed CDPK3 dependency, including AP2VIII-2 (TGGT1_233120), AP2XI-2 (TGGT1_310900), MCM7 (TGGT1_237220), and a predicted GTPase, Ras-related Rab11 (TGGT1_289680). The function of these proteins in the CDPK3 signalling cascade is less clear but may point towards biological processes related to DNA replication and transcription.

In addition to its likely involvement in the numerous signalling processes mentioned above, CDPK3 also directly or indirectly regulates the activities of other signalling proteins. For example, CDPK3-dependent down-regulation was detected at a site within the protein kinase domain of the cell-cycle-associated protein kinase GSK (TGGT1_265330 S214). Similarly, CDPK3-dependent DR^UP^ sites were found within the PI3Ka domain of PI3/4-kinase (TGGT1_215700) and the EF-hand domains of centrin 2 (TGGT1_250340) and calmodulin (TGGT1_305050).

It is important to note that visualisation of the CDPK3-dependent DR^UP^ clusters revealed that for many sites, the effect of CDPK3 deletion was temporary, such that there was an initial delay in phosphorylation, but by 60s the sites reached only slightly lower log2FC values than in WT (Fig. 3C). This may point to the redundant or compensatory activity of a protein kinase other than PKG, which could in part account for the fact that egress still occurs in CDPK3-depleted parasites, albeit at a delayed pace. Such a delay was not clearly detectable in the CDPK3-dependent DR^DOWN^ clusters.

The GO terms “signal transduction” and “hydrolase activity” mentioned above contained numerous phosphorylation sites on proteins potentially involved in cNMP signalling, including the PDEs TGGT1_202540 (PDE1) and TGGT1_241880 (PDE9), the adenylate cyclase TGGT1_270865, the guanylyl cyclase TGGT1_254370, and the CNBD containing protein TGGT1_295850 (Fig. 3E; Data S3). In addition to these previously detected targets, we also observed increased phosphorylation of PDEs TGGT1_280410 (PDE7) and TGGT1_293000 (PDE2) as well as components of phosphoinositide signalling including the PI-PLC TGGT1_248830 and the DAG kinase 1 (DGK1) TGGT1_202460. Many phosphorylation sites on these enzymes are found to be CDPK3- dependent (Fig. 3E). PI-PLC is required for the production of IP3 which triggers calcium release and has been implicated as a key downstream mediator of PKG activity (Brochet et al., 2014), while the DGK1 has been shown to play an important role in the conversion of intracellular DAG to PA and, by extension, the activation of microneme secretion and subsequent egress (Bullen et al., 2016). Collectively, these findings further substantiate our hypothesis that the A23187-mediated release of Ca^2+^ activates a CDPK3-dependent feedback loop that regulates the PKG signalling pathway.

### Deletion of CDPK3 leads to disruptions in cAMP levels and lipid signalling following A23187 treatment

While it is uncertain whether the above-mentioned phosphorylation events modulate the function of the proteins that they are found on, the putative CDPK3-dependent regulation of cNMP signalling is intriguing, as altered hydrolysis of either cAMP or cGMP has defined regulatory consequences for PKG activation. Similarly, both PI-PLC and the DAG kinase are understood to be key players in the PKG signalling pathway leading to egress (Brochet et al., 2014; Bullen et al., 2016). We therefore set out to determine whether deletion of CDPK3 would lead to disruptions in these signalling pathways.

To test whether levels of DAG and global lipid production are dysregulated following disruption of CDPK3, we performed kinetics experiments in which we analysed DAG and phospholipid levels in WT and ΔCDPK3 parasites before and after stimulus with the calcium ionophore A23187, in a similar manner to the previous phosphoproteomic timecourse. Extracellular WT and ΔCDPK3 parasites were shifted to 37°C for 60 seconds to acclimatise, and then stimulated by addition of media containing A23187 similar to established methods (Katris et al., 2020). The parasites were incubated for 5, 10, 30, 45 or 60 seconds before quenching to stop the signal chain, followed by lipid analysis. A 0s (DMSO) control was also included. After 60 seconds of A23187 stimulus, WT parasites produced slightly more DAG than ΔCDPK3 parasites, however there was no difference with the DMSO control (Fig. 4A, Supp Fig. 6A). Accordingly, WT parasites began to show less phospholipids than ΔCDPK3 parasites after 45 seconds of stimulus (Fig. 4A, Supp Fig. 6B), consistent with a lack of turnover of phospholipids to produce DAG in the ΔCDPK3 knockout parasites. While DAG-related proteins were the primary lipid related proteins affected in ΔCDPK3 based on our timecourse phosphoproteome, we also identified other proteins involved in palmitoylation and triacylglycerol synthesis that were differentially regulated, so we further investigated other lipids including Free Fatty Acids (FFAs) and triacylglycerols (TAGs). We observed a trend towards an increase in the levels of free fatty acids (FFAs) in WT parasites following A23187 stimulus (Fig. 4B, Supp Fig. 6C), which remained unchanged in ΔCDPK3 parasites. This was accompanied by a concomitant change in triacylglycerols (TAGs) whereby prior to stimulus, ΔCDPK3 parasites had more TAGs than WT parasites, but after A23187 stimulus, WT tachyzoites produced more TAGs over time so that levels became similar between both parasite lines (Fig. 4C, Supp Fig. 6D).This shows that following A23187 treatment, ΔCDPK3 parasites have altered FFA and TAG abundance necessary for lipid recycling and storage, consistent with a speculated role for CDPK3 in metabolic regulation (Treeck et al., 2014). A full lipidomic analysis of individual phospholipid species including lysophosphatidylinositol (LPI), lysophosphatidylcholine (LPC) phosphatidylinositol (PI), phosphatidylserine (PS), phosphatidylthreonine (PT) phosphatidicacid (PA), phosphatidylethanolamine (PE), FFAs, sphingomyelin (SM) and phosphatidylcholine (PC) found no significant difference in any phospholipids between WT and ΔCDPK3 parasites under normal cell culture conditions (Supp Fig. 6E), showing that defects in lipid signalling in ΔCDPK3 parasites can only be seen following A23187 calcium stimulus.

**Figure 4.**
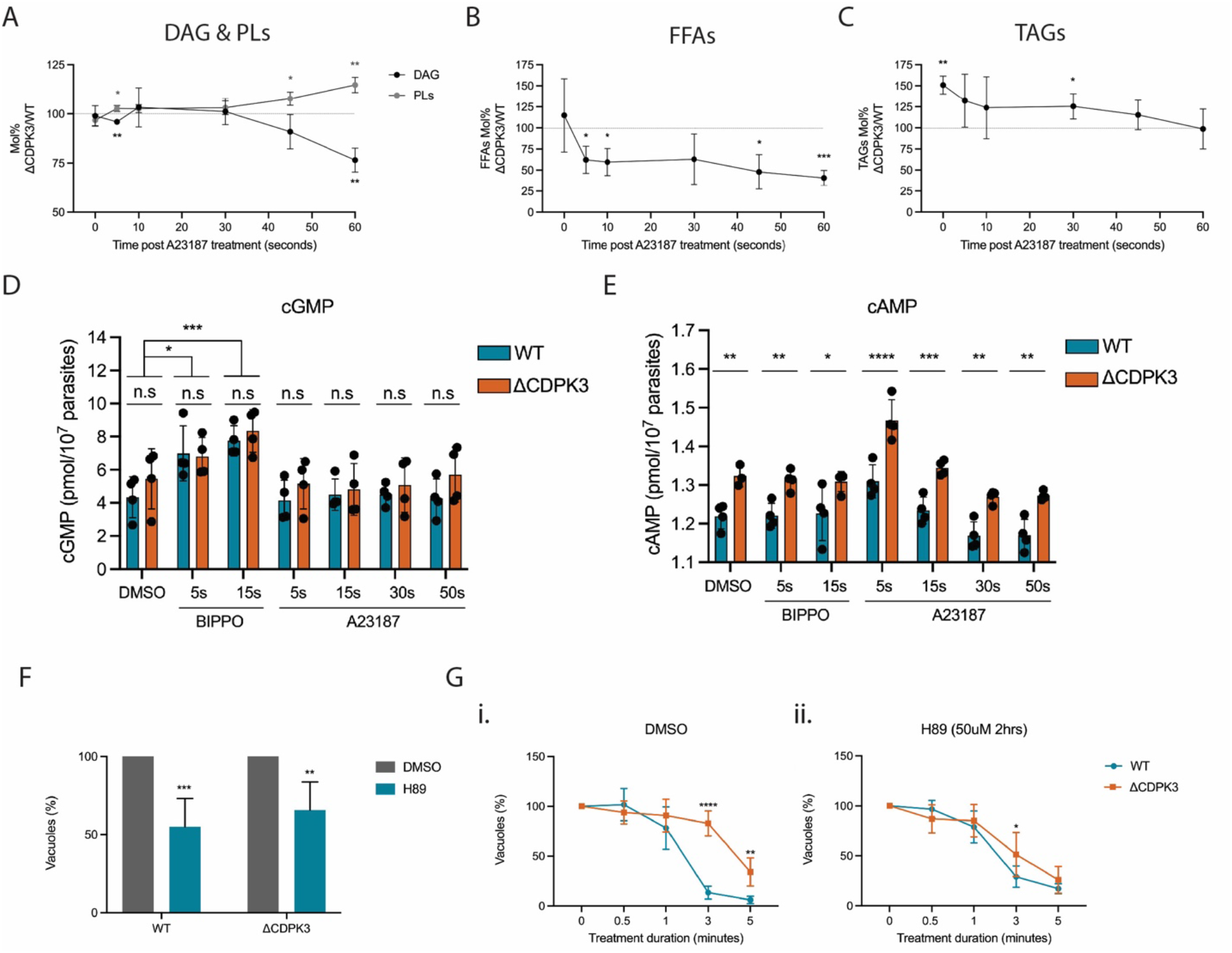
Disruption of CDPK3 leads to perturbations in lipid and cAMP signalling following A23187 treatment. Pulse experiment of WT and ΔCDPK3 parasites treated with DMSO or 8 μM A23187 for 0, 5, 10, 30, 45 or 60 seconds analysing levels of **(A)** DAG and PLs **(B)** FFAs and **(C)** TAGs, with data expressed as a ratio of ΔCDPK3/WT levels. Data are represented as mean ± s.d. (n=4). Significance was assessed using a one same t and Wilcoxon test. **, *P* ≤ 0.01; *, *P* ≤ 0.05. **(D)** Comparison of intracellular cGMP and **(D)** cAMP levels in WT and ΔCDPK3 tachyzoites following treatment with DMSO for 60 seconds; BIPPO for 5 or 15 seconds; or A23187 for 5, 15, 30 or 60 seconds. All samples were lysed in 0.1 M HCl to inactivate all PDEs, and extracts were analyzed by using commercial ELISA-based cGMP and cAMP detection assays. Data are represented as mean ± s.d. (n=4). Two-way ANOVA with Sidak multiple comparisons. ****, *P* ≤ 0.0001; ***, *P* ≤ 0.001; **, *P* ≤ 0.01; *, *P* ≤ 0.05; n.s, not significant. **(F)** Quantification of natural egress of WT and ΔCDPK3 parasites following treatment with DMSO or 50 µM H89 (2 hrs). Graph shows the remaining % of un-egressed vacuoles (relative to untreated). Data are represented as mean ± s.d. (n=5). Significance was assessed using an unpaired two-tailed t-test. ***, *P* ≤ 0.001; **, *P* ≤ 0.01 **(G)** Egress assay of **(i)** DMSO- and **(ii)** H89-pre-treated WT and ΔCDPK3 parasites following treatment with 8 µM A23187. Data are represented as mean ± s.d. (n=5). Two-way ANOVA with Holm-Sidak *post hoc* comparison. ****, *P* ≤ 0.0001; **, *P* ≤ 0.01.

Having identified several cNMP-related proteins that were differentially phosphorylated in ΔCDPK3 parasites in our phosphoproteome timecourse, we next wanted to determine whether there are any perturbations to cNMP signalling upon deletion of CDPK3 by measuring the changes in intracellular levels of cAMP and cGMP. To do this WT and ΔCDPK3 extracellular parasites that were syringe lysed in BSA free Endo buffer (Endo et al., 1987) were treated with vehicle (DMSO), 50 μM BIPPO for 5 or 15 seconds or 8 μM A23187 for 5, 15, 30 or 60 seconds at 37°C. We found that basal levels of cGMP were identical in WT and ΔCDPK3 parasites and treatment with the PDE inhibitor BIPPO resulted in elevated cGMP levels in both parasite lines compared to baseline (Fig. 4D). In contrast, and somewhat surprisingly, we found no changes in cGMP levels following A23187 treatment over the course of 60 seconds. It is important to note that our measurements of cGMP levels following treatment with A23187 differ from results shown in Stewart et al. (2017). One key difference is that we kept tachyzoites in Endo buffer, a potassium-rich buffer which mimics the intracellular environment. Stewart et al used extracellular parasites in low potassium buffer, which may have profound effects on downstream signalling pathways, given that those parasites have received signals that they are outside of the host cell. Indeed, McCoy et al have shown that the A23187- mediated egress defect of ΔCDPK3 parasites can be rescued if saponin permeabilised infected host cells are incubated in extracellular buffer, but not when incubated in intracellular buffer (McCoy et al., 2012), suggesting that there are key signalling differences between intra and extracellular parasites.

Basal cAMP levels, by contrast, were 8.3% higher in knockout parasites compared to WT. Following treatment with A23187, we found that cAMP levels initially rose in both WT and knockout parasites at 5 seconds post treatment, with a gradual decrease to below basal levels at 60 seconds post treatment (Fig. 4E). We observed no immediate change in cAMP levels following BIPPO treatment, suggesting that BIPPO does not inhibit cAMP-specific PDEs in *T. gondii*. These findings point towards perturbations in cAMP signalling in ΔCDPK3 parasites which have elevated basal levels that further increase upon A23187 treatment.

Since basal cAMP levels are elevated in ΔCDPK3 parasites compared to WT, we reasoned that inhibition of cAMP signalling would overcome the A23187-mediated egress delay observed in knockout parasites. To test this, we treated both WT and ΔCDPK3 parasites with the ATP-competitive PKAc inhibitor H89 (50 µM). After 2 hours of treatment, there was a significant amount of premature egress in both WT and ΔCDPK3 parasites (∼45% & ∼34%, respectively; Fig. 4F) consistent with previous reports that have shown that the downregulation of cAMP signalling by genetic disruption of PKAc1 stimulates premature egress in *Toxoplasma* (Jia et al., 2017; Uboldi et al., 2018). Intriguingly, when we investigated A23187-induced egress rates of the remaining intracellular H89 pre-treated parasites, we found that the egress delay normally observed in ΔCDPK3 parasites was largely rescued with H89 pre-treatment (Fig. 4Gi-ii). This finding suggests that pharmacological inhibition of cAMP signalling is sufficient to partially compensate for the deletion of CDPK3.

### Cell biological and biochemical characterisation of PDE1, 2, 7 and 9

The preponderance of A23187-induced phosphorylation on several PDEs suggests that they may play an important role in the cAMP- and cGMP-mediated signalling cascades that lead to egress assuming that phosphorylation may directly, or indirectly control their activity. Specifically, we predicted a cAMP-specific PDE would play an important role given the rescue of the egress defect observed in ΔCDPK3 parasites by the PKA inhibitor H89. As the specificity for the majority of *Toxoplasma* PDEs has not been experimentally validated, we generated HA-tagged conditional knockouts (cKOs) of the 4 PDEs identified as being phosphorylated following A23187 treatment in order to characterise them and identify which are capable of hydrolysing cAMP (Fig. 5A). For each line, integration of both repair templates was validated by PCR (Supp Fig. 7). Western blot analysis confirmed that they migrate at their predicted sizes (Fig. 5B), and we found that each PDE occupies a distinct cellular localisation (Fig. 5C), in agreement with a previous report (Vo et al., 2020; Moss et al., 2021). To identify the substrate specificity of the PDEs, we immunoprecipitated each via the HA-tag from parasite lysates and measured their hydrolytic activity *in vitro*. We included the *P. falciparum* PDEβ that was previously shown to be dual-specific as a positive control (Flueck et al., 2019). PDEs 1, 7 and 9 were able to hydrolyse cGMP, while PDE2 is specific for cAMP (Fig. 5D). Only PfPDEβ displayed dual-hydrolytic activity in our hands.

**Figure 5.**
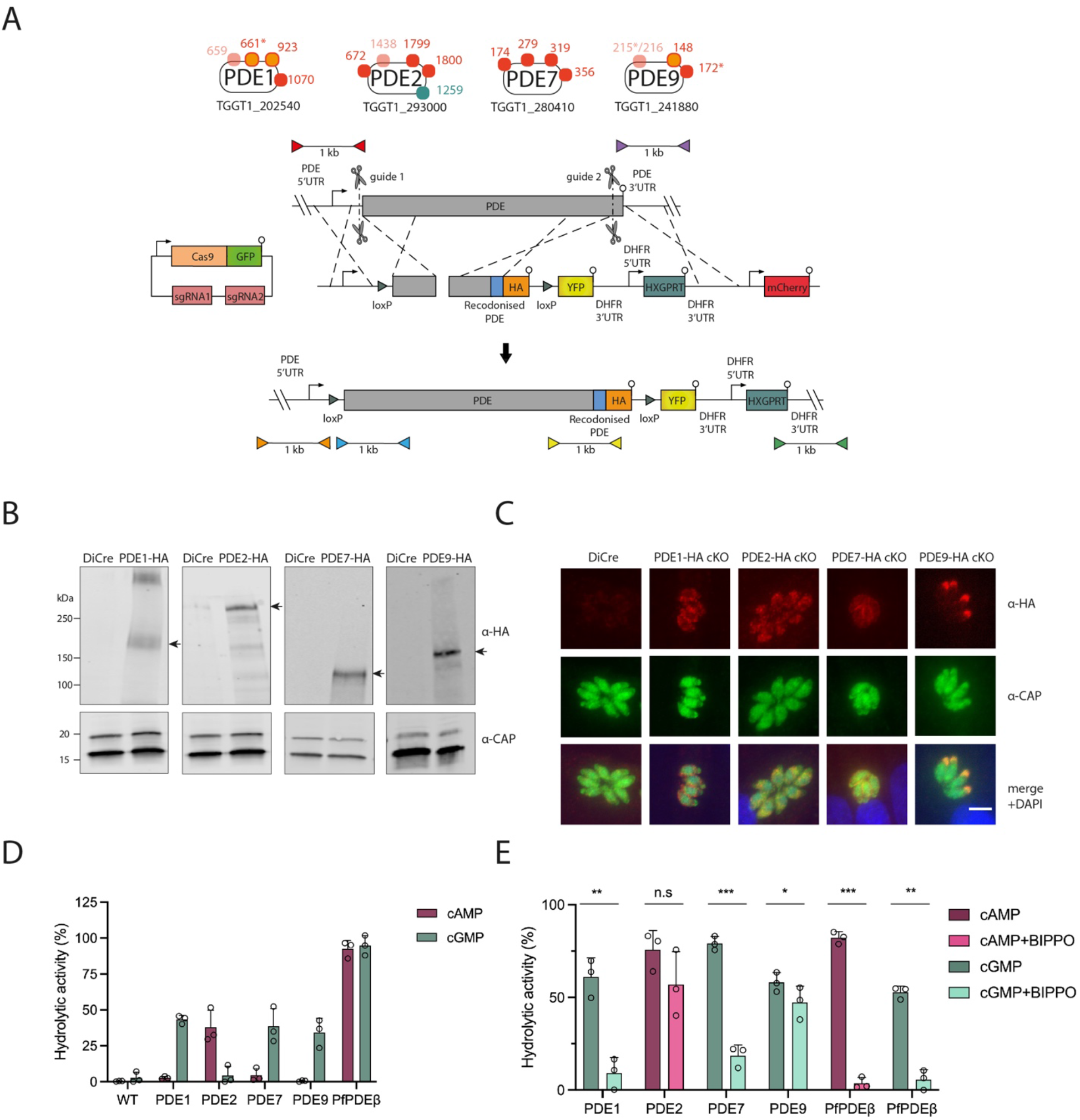
Candidate PDEs occupy distinct cellular localisations and are differentially inhibited by BIPPO. **(A)** Schematic representation of the PDEs identified in the timecourse phosphoproteome (see Fig. 2E) and the strategy use to generate the conditional PDE knockout lines. CRISPR/Cas9 was used to generate two cuts in the gene and two separate repair templates were provided to integrate one loxP site (green triangle) upstream of the PDE gene, and another repair template to tag the PDE with a C-terminal HA epitope tag (orange) and introduce a second loxP site, a YFP sequence and the HXGPRT cassette. Scissors represent Cas9 cleavage sites and lollipops depict stop codons. Coloured triangles represent primer pairs used to detect WT, 5’ integration and 3’ integration loci for 5’ loxP integration (red, orange and blue respectively) and 3’ tagging (purple, yellow and green respectively). PCR results using these primer pairs are shown in Supp Fig. 7. **(B)** Western blot analysis of parental DiCre and HA-tagged PDE1, PDE2, PDE7 and PDE9 cKO parasites probed with α-HA antibodies showing migration of the PDEs at their expected molecular weights as depicted by arrows. A non-specific band >250 kDa is observed in the PDE1- HA cKO line. Blots were probed with α-CAP antibodies as a loading control. **(C)** Immunofluorescence analysis of DiCre and HA-tagged PDE1, PDE2, PDE7 and PDE9 cKO lines probing with α-HA (red) and α- CAP (green) antibodies. Scale bar, 5 μm. **(D)**Hydrolytic activity of immunoprecipitated HA-tagged PDE1, PDE2, PDE7, PDE9 and the PfPDEβ using either cAMP or cGMP as a substrate. Lysates from the WT parental line were also included as a control. Data are represented as mean ± s.d. (n=3). **(E)** Hydrolytic activity of immunoprecipitated HA-tagged PDE1, PDE2, PDE7, PDE9 and PfPDEβ after incubating with DMSO (vehicle) or 25 μM BIPPO. cAMP was used as a substrate for PDE2 and PDEβ, while cGMP was used as a substrate for PDE1, PDE7, PDE9 and PDEβ. Data are represented as mean ± s.d. (n=3). Significance was assessed using a paired t-test. ***, *P* ≤ 0.001; **, *P* ≤ 0.01; *, *P* ≤ 0.05; n.s. not significant.

To further confirm the hydrolytic specificity of the PDEs, we treated each of the samples with BIPPO. While the specificity of BIPPO has not been experimentally validated in *Toxoplasma*, cGMP-specific PDE1 and 7 were significantly inhibited by BIPPO, while the cAMP-specific PDE2 was refractory to BIPPO inhibition (Fig. 5E). Interestingly, PDE9, a cGMP-specific PDE appears less sensitive to BIPPO treatment. This is in agreement with a previous study (Vo et al., 2020), although in our hands PDE9 is cGMP-specific and not dual-specific. Collectively these data show that BIPPO is a cGMP-specific inhibitor in *Toxoplasma* and lends further support that PDE2 is a cAMP-specific PDE. Interestingly, we found that both the cAMP- and cGMP-hydrolysis activities of PfPDEβ are inhibited with BIPPO. It will be interesting in the future to evaluate the structural differences between the PDEs and the inhibitory potential of BIPPO.

### Functional assessment of the candidate PDEs reveals that PDEs 1 & 2 are important but not essential during the lytic cycle while PDEs 7 & 9 are dispensable

We next wanted to establish which of the aforementioned PDEs were essential for lytic growth. Addition of rapamycin (RAP) to the HA-tagged PDE cKO lines leads to excision of the PDE gene of interest in the respective cKO lines (Fig. 6A). Despite near complete excision in all lines as observed by PCR 24 hours post RAP treatment (Fig. 6B), it was only until 3 days post-treatment that we saw complete protein depletion below detectable levels (Fig. 6C-D). Therefore, all subsequent experiments were conducted with parasites 3 days post RAP treatment. To assess the impact of PDE disruption on parasite viability and growth, we performed plaque assays using DMSO- and RAP-treated PDE cKO parasites and measured the size and number of plaques after 5 days (Fig. 6E-G). All knockout lines were able to form plaques, with deletion of PDE7 and PDE9 resulting in no significant changes in plaque size or number. However, PDE1 and PDE2 formed much smaller plaques, with a 37% and 81% reduction in plaque sizes respectively. Despite a marked reduction in plaque size for PDE2 knockout parasites, there was no significant change in the number of plaques compared to the DMSO-treated line. Deletion of PDE1 on the other hand resulted in a 4-fold reduction in the number of plaques formed. Overall, these results suggest that both PDE1 & PDE2 are important but not essential for lytic growth and that PDE1 may be important for egress and/or invasion due to the reduced number of plaques formed following its disruption. Reassuringly, our observed phenotypes for PDE 1, 2, 7 and 9 KOs are in agreement with a recent study which knocked down all predicted *Toxoplasma* PDEs (Moss et al., 2021).

**Figure 6.**
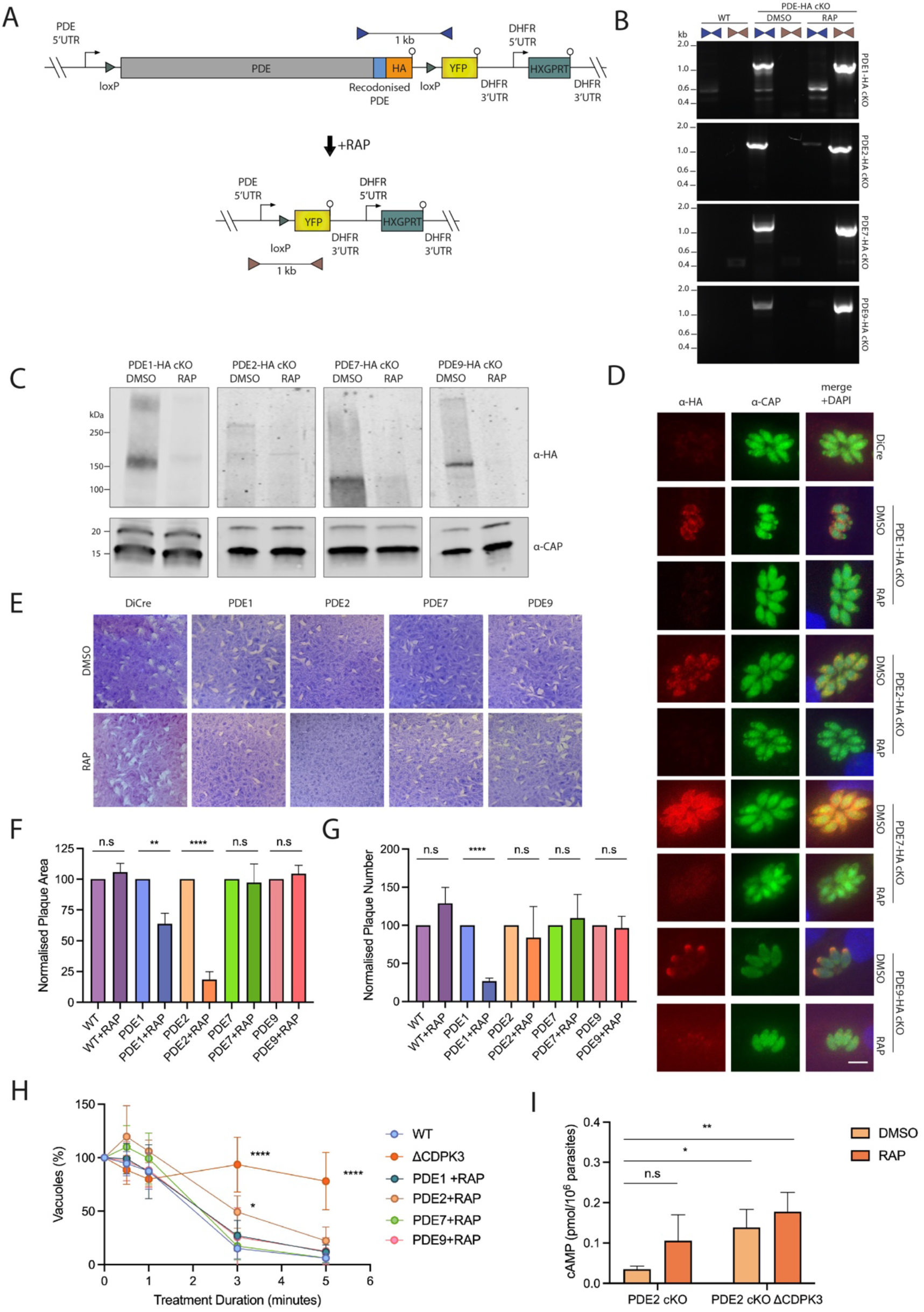
PDE1 and PDE2 are important for lytic growth, with ΔPDE2 parasites displaying an A23187- mediated egress defect similar to ΔCDPK3 parasites. **(A)** Schematic representation of PDE2 rapamycin mediated deletion of PDE cKO lines. Addition of rapamycin leads to excision of the entire gene, placing YFP under the control of the PDE promoter. Coloured triangles represent primer pairs used to detect unexcised (blue) and excised (brown) loci. **(B)** PCR analysis of DMSO- and RAP-treated PDE cKO parasite lines showing near complete excision for all lines. **(C)** Western blot analysis of PDE cKO lines showing near complete loss of the PDEs at the protein level following treatment with RAP **(E)** Representative images of plaque assays performed on DMSO- and RAP-treated WT and PDE cKO lines after a period of 5 days. Assays were performed in biological triplicates. **(F)**Measurement of plaque area shown in Fig. 6E. Data are represented as mean ± s.d. (n=3). Significance was assessed using multiple t-tests. ****, *P* ≤ 0.0001; **, *P* ≤ 0.01; n.s, not significant. **(G)** Quantification of plaque numbers shown in Fig. 6E. Data are represented as mean ± s.d. (n=3). Significance was assessed using multiple t-tests. ****, *P* ≤ 0.0001; n.s. not significant. **(H)** Egress assay of GFP-T2A-jRCaMP1b expressing WT, ΔCDPK3 and RAP-treated PDE cKO parasites following treatment with 8 µM A23187. Data are represented as mean ± s.d. (n=3). Two-way ANOVA.****, *P* ≤ 0.0001; *, *P* ≤ 0.05. **(I)** Quantification of intracellular cAMP levels of PDE2 cKO and PDE2 cKO ΔCDPK3 parasites treated with either DMSO or RAP. Data are represented as mean ± s.d. (n=4). Two-way ANOVA with Turkey’s multiple comparisons. **, *P* ≤ 0.01; *, *P* ≤ 0.05.

We next wanted to determine whether disruption of any of the PDEs would lead to an A23187-mediated egress delay, similar to ΔCDPK3 parasites. We reasoned that if one of the PDEs is involved in the A23187-mediated feedback loop, then disruption of this PDE would mimic, at least partially, the A23187-mediated egress delay observed in ΔCDPK3 parasites. Using a medium-throughput plate-based egress assay we found that only deletion of PDE2 showed a modest egress defect (Fig. 6H, Supp Fig. 8). This suggests an important, but non-essential role for PDE2 in cAMP signalling. Since the egress defect observed upon deletion of PDE2 did not reach the severity of the egress defect observed in ΔCDPK3 parasites, it is likely that other PDEs, and/or cyclases are involved in the dysregulation of cAMP levels found in our study, some of which may have been missed in our phosphoproteome or may not be regulated by phosphorylation. To test this, we generated a ΔCDPK3 KO in the PDE2 cKO line using a similar approach to the one used in Fig. 2A, however we substituted the HXGPRT cassette with a DHFR-TS selection cassette. Upon deletion of PDE2, we observed a further increase of cAMP levels compared to either single gene deletion (Fig. 6I) suggesting that CDPK3 is regulating another unknown cAMP-specific PDE, or an adenylyl cyclase.

## Discussion

In this study we have aimed to unravel the complexity of the signalling pathways that govern the control of host cell egress of *Toxoplasma* from its host cell. Several signalling components conserved in higher eukaryotes have previously been identified, and their connectivity, to some extent described. However, the published data is not currently supported by a model that fits most experimental results. A deeper understanding of how the signalling pathways are interconnected is essential for our understanding of the regulation of host cell egress and of the integration of environmental or endogenous signals in general. This is important since we do not fully understand how *Toxoplasma* parasites sense and react to their environment. Furthermore, the plethora of calcium- dependent kinases and phosphodiesterases imply a highly complex and sensitive interplay of signalling pathways to finetune cellular responses to inputs. Our results are likely of importance beyond *Toxoplasma gondii* research: the plant-like calcium- dependent kinases (CDPKs) are conserved in *Plasmodium* species where it has been shown that PDE inhibitors can overcome disruption of CDPKs (Absalon et al., 2018).

Our data quite clearly support previous data on the importance of cNMP, Ca^2+^ and lipid signalling on parasite exit from the host cell. They furthermore add an important new layer of information: CDPK3, and potentially other kinases that depend on CDPK3 function, are part of a feedback loop that enables rapid signalling. This feedback loop does not control release of Ca^2+^ from internal stores, since McCoy *et al* (2012) have shown that disruption of CDPK3 does not lead to a delay in Ca^2+^ release. It also does not appear to act through elevation of cGMP levels as we show both that treatment with ionophore does not lead to a measurable increase in cGMP levels and that cGMP levels are not dependent on CDPK3.

More likely, and in keeping with our cAMP measurements, CDPK3 directly or indirectly downregulates levels of cAMP. This, in turn, alters activity of the cAMP-dependent protein kinase, PKAc. It is interesting to note that Jia and colleagues found a clear dependency on PKG for parasites to egress upon PKAc depletion, but they were equally unable to reliably ascertain cGMP accumulation in intracellular parasites (Jia et al., 2017). While it is possible that our collective inability to observe elevated cGMP levels is explained by the sensitivity limits of the assay employed, it is similarly possible that cAMP-mediated signalling is exerting its effects on the PKG signalling pathway in a cGMP-independent manner. While no such mechanism has been described, it is possible that phosphorylation of PKG may lead to changes in its affinity for cGMP or it may regulate the activity of the kinase itself. Further work is needed to clarify the role of cGMP levels in these conditions. We also identified dysregulation of DAG and phospholipid signalling in ΔCDPK3 parasites following A23187 treatment, which could be contributing to the delayed egress phenotype observed in the KO parasites. Having identified CDPK3- dependent phosphorylation sites on both DGK1 and PI-PLC in our timecourse phosphoproteome, it is possible that these perturbations are being directly mediated by CDPK3. Alternatively, and as outlined above, any changes in cGMP levels or PKG activity in ΔCDPK3 parasites could also lead to the dysregulation of phospholipid signalling we observed.

We identify PDE2 as one contributor of cAMP control, however, through double gene deletions of PDE2 and CDPK3, we show that other cAMP signalling components likely contribute to a further cAMP imbalance. This could be either via as yet unidentified PDEs with cAMP specificity or, more likely, an adenylate cyclase. In support of the latter, we identified several CDPK3-dependent phosphorylation sites on an ACβ following A23187 treatment. We also found that deletion of PDE2 alone leads to a modest egress phenotype that does not reach ΔCDPK3 levels. This is not surprising: our data, and previous studies have identified many CDPK3-dependent targets (Treeck et al., 2014; Gaji et al., 2015; McCoy et al., 2017; Wallbank et al., 2019), and the CDPK3-mediated phenotype is likely caused by a combination of the phosphorylation events identified here. However, we did not detect any CDPK3-dependent phosphosites on PDE2, so a direct link between CDPK3 and PDE2 is currently missing. However, it is possible that CDPK3- dependent phosphorylation sites on PDE2 were not detected in our mass-spectrometry experiments for technical reasons, or that PDE2 is indirectly regulated by CDPK3. It has, for instance, been reported that CDPK3 promotes egress by phosphorylating the egress suppressor SCE1 (McCoy et al., 2017). Deletion of SCE1 in ΔCDPK3 parasites largely rescues several CDPK3-dependent phosphosites, suggesting that another SCE1- suppressed kinase is able to partially compensate for loss of CDPK3, and likely amplifies regular CDPK3-mediated egress under normal conditions.

The research community is also continuously identifying novel components involved in signalling which, once identified, could shed light on how different pathways are interconnected. For example, a recent report has identified SPARK, a novel kinase that appears to mediate Ca^2+^ release in a PKG-dependent manner and can be largely bypassed via treatment with A23187 (Smith et al., 2021). While A23187 treatment appears to restore absolute levels of Ca^2+^ release in SPARK depleted parasites, the rate of both calcium release and egress remains partially delayed. These findings suggest that PKG-regulated SPARK still contributes, to some degree, to A23187-mediated egress. This observation is in keeping with our proposition that A23187 signalling feeds back into the PKG signalling pathway.

While our study provides strong evidence for a CDPK3-mediated feedback loop to control rapid egress and nearly overlapping signalling pathways at peak calcium flux, we cannot draw conclusions about the signalling events at the onset of calcium release. Nevertheless, our timecourse data identifies rapid CDPK3-dependent differences on proteins involved in cNTD signalling as early as 15 seconds post-induction, suggesting that it plays a role at the very onset of the signalling cascades, well before calcium release peaks. We are still far from identifying all players in the signalling cascades that lead to egress from the host cell. Arrayed CRISPR screens targeting of the Toxoplasma kinome in ΔCDPK3 parasites (Young et al., 2019; Smith et al., 2021) will likely shed further light on these signalling pathways.

## Materials and Methods

### Parasite culture and transfection

*T. gondii* tachyzoite RH strains lacking KU80 (*Δku80*) and HXGPRT (Δ*hxgprt*) (Fox et al., 2009; Huynh and Carruthers, 2009) were cultured in a confluent monolayer of human foreskin fibroblasts (HFFs) maintained in Dulbecco’s Modified Eagle medium GlutaMAX™(DMEM+ GlutaMAX™, Gibco) supplemented with 10% foetal bovine serum (FBS), at 37°C and 5% CO2.

### Plasmid and parasite strain generation

Primers used throughout this study are listed in Table S6. The calcium sensor construct was generated as recently described (Alves et al., 2021). The construct was linearised using NaeI and transfected into RH *Δku80Δhxgprt* parasites as described previously (Soldati and Boothroyd, 1993) to generate the GFP-T2A-jRCaMP1b calcium sensor line. Transgenic parasites were subjected to 5’-fluo-2’-deoxyuridine (FUDR) selection (5 µM) 24 hrs after transfection. To generate the GFP-T2A-jRCaMP1b ΔCDPK3 line, the *HXGPRT* casette (flanked by 5’ and 3’ DHFR UTR sequences) was PCR amplified from *pGRA*-HA_HXGPRT (Coppens et al., 2006) using primers 1/2 (introducing 40bp CDPK3 homology regions to the amplified fragment) and co-transfected into RH *Δku80Δhxgprt* with pSag1::Cas9-U6::dbl-sgCDPK3. The pSag1::Cas9-U6::dbl-sgCDPK3 vector was generated by inverse PCR amplification of the pSag1::Cas9-U6 (Behnke et al., 2014) vector using primer pairs 3/4 and 3/5 to generate intermediate constructs pSag1::Cas9- U6::sg1CDPK3 (comprising sgRNA1) and pSag1::Cas9-U6::sg2CDPK3 (comprising sgRNA2) respectively. Following circularization of both intermediate constructs using KLD reaction buffer (NEB), a region comprising sgRNA1 was PCR amplified with primers 6 and 7 from pSag1::Cas9-U6::sg1CDPK3 and Gibson assembled into Kpn1/XhoI linearised pSag1::Cas9-U6:: sg2CDPK3 to generate the double sgRNA plasmid pSag1::Cas9-U6::dbl-sgCDPK3. Recombinant parasites were selected 24 hrs post transfection by addition of mycophenolic acid (MPA; 25µg/mL) and xanthine (XAN; 50 µg/mL) to culture medium. Integration of the HXGPRT cassette at the CDPK3 locus was confirmed using primer pairs 8/9 and 10/11 to confirm 5’ and 3’ integration respectively. Absence of the endogenous CDPK3 locus was confirmed using primers 12/13.

To generate the PDE1, PDE2, PDE7 and PDE9 HA-tagged conditional knockout lines, two separate repair templates were generated for each gene; one which would integrate a loxP site 100 bp upstream of the start codon, and one that would introduce a C-terminal HA epitope tag along with a second loxP site and an HXGPRT cassette downstream of the gene.

To generate the pUC19_PDE1_5’loxP repair construct, a 1 kb 5’ homology region and a 1 kb 3’ homology region were PCR amplified from genomic DNA using primers 14/15 and 16/17 respectively, with the primers designed to introduce a loxP site between the 5’ and 3’ homology regions. The fragments were then Gibson cloned into the BamHI and EcoRI sites of the pUC19 vector. To generate the pG140_PDE1-HA_3’loxP_HXGPRT plasmid, a 1 kb 5’ homology region was amplified from genomic DNA using primers 18/19 and the HA tag was amplified from an unpublished in-house plasmid using primers 20/21. These fragments were Gibson cloned into the HindIII & PacI sites of the pG140 plasmid to generate an intermediate plasmid. A 1 kb 3’ homology region was PCR amplified from genomic DNA using primers 22/23, while an mCherry coding sequence flanked by Gra gene UTRs was amplified from pTKO2C (Caffaro et al., 2013) using primers 24/25. These fragments were subsequently Gibson cloned into the SacI sites of the intermediate plasmid to generate pG140_PDE1-HA_3’loxP_HXGPRT.

The pSag1::Cas9-U6::dbl-sgPDE1 vector was generated by inverse PCR amplification of the pSag1::Cas9-U6 (Behnke et al., 2014) vector using primer pairs 3/26 and 3/27 to generate intermediate constructs pSag1::Cas9-U6::sg1PDE1 (comprising sgRNA1) and pSag1::Cas9-U6::sg2PDE1 (comprising sgRNA2) respectively. Following circularization of both intermediate constructs using KLD reaction buffer (NEB), a region comprising sgRNA1 was PCR amplified with primers 6 and 7 from pSag1::Cas9-U6::sg1PDE1 and Gibson assembled into Kpn1/XhoI linearised pSag1::Cas9-U6:: sg2PDE1 to generate the double sgRNA plasmid pSag1::Cas9-U6::dbl-sgPDE1.

After linearising pUC19_PDE1_5’loxP with HindIII & EcoRI and pG140_PDE1- HA_3’loxP_HXGPRT with HindIII & SapI, the two repair templates were co-transfected with pSag1::Cas9-U6::dbl-sgPDE1 into the RH DiCre *Δku80Δhxgprt* line (Hunt et al., 2019). Recombinant parasites were selected 24 hrs post transfection by addition of mycophenolic acid (MPA; 25µg/mL) and xanthine (XAN; 50 µg/mL) to culture medium.

The same cloning strategy was used for all other PDE cKO lines with the primer pairs used in each step listed in table S1.

To generate the PDE2-HA-cKOΔCDPK3 line, the *DHFR-TS* casette (flanked by GRA1 5’ and GRA2 3’ UTR sequences) was PCR amplified from an unpublished in-house plasmid using primers 28/29 (introducing 40bp CDPK3 homology regions to the amplified fragment) and co-transfected into RH *Δku80Δhxgprt* with pSag1::Cas9-U6::dbl- sgCDPK3. Recombinant parasites were selected 24 hrs post transfection by addition of pyrimethamine (1 µM) to culture medium.

### Egress assay

Fresh tachyzoites were harvested and seeded onto confluent HFF monolayers in black 96-well imaging μ-plates (Ibidi) at an MOI of 0.5. After 28 hours of growth, egress assays were performed in triplicate at 37 °C in Ringers buffer (155 mM NaCl, 3 mM KCl, 2 mM CaCl2, 1 mM 556 MgCl2, 3 mM NaH2PO4, 10 mM HEPES, 10 mM glucose). The parasites were incubated with 8 µM Ca2+ ionophore A23187 (BioVision) or 50 µM BIPPO (generated in-house) for variable timings. Wells were subsequently fixed by adding 16% FA to a final concentration of 3% for 15 mins. Wells were washed with PBS and stained with 5 µg/ml DAPI. Automated image acquisition of 25 fields per well was performed on a Cellomics 561 Array Scan VTI HCS reader (Thermo Scientific) using a 20× objective. Image analysis was performed using the Compartmental Analysis BioApplication on HCS Studio (Thermo Scientific).

### Live imaging of calcium sensor line

Fresh tachyzoites were harvested and seeded (at an MOI of 0.5) onto confluent HFF cells grown on IBIDI tissue culture treated 8 well chamber slides and allowed to grow for 28 hrs in DMEM + 10%FBS. Prior to imaging, wells were washed once with PBS, and supplemented with 100 µl Ringer’s media. Wells were treated for 5 mins at 37°C with 100 µl 2 µg/ml Cytochalasin D in Ringer’s buffer (final concentration 1 µg/ml) to prevent egress. Imaging was performed on the Nikon Eclipse Ti-U inverted fluorescent microscope, 60x/1.4 NA Oil immersion objective, in environmental chamber (OKOLAB) with temperature maintained at 37°C. Image capture was managed by Nikon NIS-Elements software with acquisition 1/s for 70s. At 15s following image acquisition, 100µl of A23187 (24µM) or BIPPO (150 µM) in Ringer’s buffer was added by pipette (to final concentrations of 8 µM and 50 µM respectively). ≥10 vacuoles across ≥10 wells were imaged across ≥ 7 days for each condition. Image analysis was performed using Nikon NIS-Elements analysis software. jRCaMP1b and GFP signals at 0s were set to 0 (zero) and 1 respectively. jRCaMP1b/GFP was used as a readout for ΔCa^2+^.

### Phosphoproteome analysis

#### Lysis and protein digestion

Parasites were seeded onto HFF monolayers in 15cm culture dishes at an MOI of 5. 24 hours post-inoculation, plates were washed once with PBS and treated with 50 µM BIPPO (15s) or 8 µM A23187 (variable timings depending on experiment) in Ringer’s buffer. Following the appropriate treatment duration, treatments were rapidly removed and plates placed on a supercooled salt water ice bath to inhibit further signalling. Lysis was performed by scraping cells in ice cold 8 M urea, 75 mM NaCl, 50 mM Tris, pH 8.2, supplemented with protease (complete mini, Roche) and phosphatase (PhosSTOP, Roche) inhibitors. Lysis was followed by sonication to reduce sample viscosity (30% duty cycle, 3 x 30 seconds bursts, on ice). Protein concentration was measured using a BCA protein assay kit (Pierce). Lysates (1mg each) were subsequently reduced with 5 mM DTT for 30 minutes at 56 ⁰C and alkylated in the dark with 14 mM iodoacetamide for 30 minutes at RT. Following iodoacetamide quenching with 5 mM DTT for 15 minutes in the dark, lysates were diluted with 50 mM ammonium bicarbonate to < 4M urea, and digested with LysC (Promega) for 2-3 hours at 37 ⁰C. Lysates were further diluted with 50 mM ammonium bicarbonate to < 2M urea and digested with trypsin (Promega) overnight at 37 ⁰C. After digestion, samples were acidified with trifluoroacetic acid (TFA) (Thermo Fisher Scientific) to a final concentration of 1% (v/v). All insoluble material was removed by centrifugation and the supernatant was desalted on Sep-Pak C18 cartridges (Waters).

#### TMT labelling

Samples were dissolved at 1 mg/ml in 50 mM Na-Hepes, pH 8.5 and 30% acetonitrile (v/v) and labelled with respective TMT reagents (Thermo Fisher Scientific, 2.4 mg reagent/1 mg sample) for 1 hour at RT. Labelling was then quenched with 0.3% hydroxylamine for 15 minutes at RT and samples acidified (pH∼2) with formic acid. After verification of labelling efficiency via mass spectrometry, the lysates were mixed in a 1:1 ratio, vacuum dried and desalted on Sep-Pak C18 cartridges.

#### Phosphopeptide enrichment

Desalted and vacuum dried samples were solubilised in 1 ml of loading buffer (80% acetonitrile, 5% TFA, 1 M glycolic acid) and mixed with 5 mg of TiO2 beads (Titansphere, 5 µm GL Sciences Japan). Samples were incubated for 10 minutes with agitation, followed by a 1 minute 2000 × g spin to pellet the beads. The supernatant was removed and used for a second round of enrichment as explained below. Beads were washed with 150 μl loading buffer followed by two additional washes, the first with 150 μl 80% acetonitrile, 1% TFA and the second with 150 μl 10% acetonitrile, 0.2% TFA. After each wash, beads were pelleted by centrifugation (1 minute at 2000 × g) and the supernatant discarded. Beads were dried in a vacuum centrifuge for 30 minutes followed by two elution steps at high pH. For the first elution step, beads were mixed with 100 μl of 1% ammonium hydroxide (v/v) and for the second elution step with 100 µl of 5% ammonium hydroxide (v/v). Each time beads were incubated for 10 minutes with agitation and pelleted at 2000 × g for 1 minute. The two elutions were removed following each spin, and subsequently pooled together before undergoing vacuum drying. The supernatant from the TiO2 enrichment was desalted on Sep-Pak C18 and the High Select Fe-NTA phosphopeptide enrichment kit (Thermo Fisher Scientific) was used according to manufacturer’s instructions for a second round of enrichment.

#### Sample fractionation and desalting

Combined TiO2 and Fe-NTA phosphopeptide eluates were fractionated using the Pierce High pH Reversed-Phase kit (Thermo Fisher Scientific) according to manufacturer’s instructions. Resulting fractions were taken to dryness by vacuum centrifugation and further desalted on a stage tip using Empore C18 discs (3M). Briefly, each stage tip was packed with one C18 disc, conditioned with 100 µl of 100% methanol, followed by 200 µl of 1% TFA. The sample was loaded in 100 μl of 1% TFA, washed 3 times with 200 µl of 1% TFA and eluted with 50 µl of 50% acetonitrile, 5% TFA. The desalted peptides were vacuum dried in preparation for LC-MS/MS analysis.

#### LC-MS/MS

Samples were resuspended in 0.1% TFA and loaded on a 50 cm Easy Spray PepMap column (75 μm inner diameter, 2 μm particle size, Thermo Fisher Scientific) equipped with an integrated electrospray emitter. Reverse phase chromatography was performed using the RSLC nano U3000 (Thermo Fisher Scientific) with a binary buffer system (solvent A: 0.1% formic acid, 5% DMSO; solvent B: 80% acetonitrile, 0.1% formic acid, 5% DMSO) at a flow rate of 250 nl/minute. The samples were run on a linear gradient of 5-60% B in 150 minutes with a total run time of 180 minutes including column conditioning. The nanoLC was coupled to an Orbitrap Fusion Lumos mass spectrometer using an EasySpray nano source (Thermo Fisher Scientific). The Orbitrap Fusion Lumos was operated in data-dependent mode using two acquisition methods. For the MS2 method, HCD MS/MS scans (R=50,000) were acquired after an MS1 survey scan (R=120, 000) using MS1 target of 4E5 ions, and MS2 target of 2E5 ions. The number of precursor ions selected for fragmentation was determined by the “Top Speed” acquisition algorithm with a cycle time of 3 seconds, and a dynamic exclusion of 60 seconds. The maximum ion injection time utilised for MS2 scans was 86 ms and the HCD collision energy was set at 38. For the MS3 method, CID MS/MS scans (R=30,000) were acquired after an MS1 survey scan with parameters as above. The MS2 ion target was set at 5E4 with multistage activation of the neutral loss (H3PO4) enabled. The maximum ion injection time utilised for MS2 scans was 80 ms and the CID collision energy was set at 35. HCD MS3 scan (R=60,000) was performed with synchronous precursor selection enabled to include up to 5 MS2 fragment ions. The ion target was 1E5, maximum ion injection time was 105 ms and the HCD collision energy was set at 65. Acquired raw data files were processed with MaxQuant (Cox and Mann, 2008; Cox et al., 2011) (version 1.5.2.8) and peptides were identified from the MS/MS spectra searched against *Toxoplasma gondii* (combined TG1, ME48 and VEG proteomes, ToxoDB) and *Homo sapiens* (UniProt, 2018) proteomes using Andromeda (CITE Cox et al. 2011) search engine. TMT based experiments in

MaxQuant were performed using the ‘reporter ion MS2 or MS3’ built-in quantification algorithm with reporter mass tolerance set to 0.003 Da. Cysteine carbamidomethylation was selected as a fixed modification. Methionine oxidation, acetylation of protein N-terminus, deamidation (NQ) and phosphorylation (S, T, Y) were selected as variable modifications. The enzyme specificity was set to trypsin with a maximum of 2 missed cleavages. The precursor mass tolerance was set to 20 ppm for the first search (used for mass re-calibration) and to 4.5 ppm for the main search. The datasets were filtered on posterior error probability to achieve a 1% false discovery rate on protein, peptide and site level. ‘Match between runs’ option was enabled for fractionated samples (time window 0.7 min) and “Unique and razor peptides” mode was selected to allow identification and quantification of proteins in groups (razor peptides are uniquely assigned to protein groups and not to individual proteins). All mass spectrometry acquisition files and MaxQuant processing files have been deposited to the ProteomeXchange Consortium via the PRIDE (Perez-Riverol et al., 2019) partner repository (currently awaiting dataset identifier).

### Phosphoproteome data processing

#### A23187/BIPPO analysis (set1 and set2)

The data were analyzed using Perseus (Tyanova et al., 2016) (version 1.5.0.9) and Microsoft Office Excel 2016. Briefly, the data were filtered to remove common contaminants, IDs originating from reverse decoy sequences and sites originating from the host (human) proteome. Individual TMT reporter intensities (MS2-based acquisition) and total intensity were log2 and log10 transformed, respectively. Log2 reporter intensities for each sample were subsequently normalised (centered) by subtracting the median log2 reporter intensity value calculated for all non-phosphorylated peptides detected in the same sample. Data were then filtered by 1 valid value to retain only the quantified phosphosites and log2 fold changes in reporter intensity between conditions were calculated. Differentially regulated (DR) phosphorylation sites were identified by calculating the median absolute deviation (MAD) for the log2FC in each comparative dataset. The largest of these was used to set an outlier threshold of 3x MAD (rounded to the nearest tenth; log2FC>0.5 for up-regulated sites and log2FC<-0.5 for down-regulated sites) and applied across all datasets.

#### A23187 timecourse analysis

The data were analyzed using Perseus (Tyanova et al., 2016) (version 1.5.0.9) and Microsoft Office Excel 2016. TMT reporter intensities obtained via MS2 and MS3-based acquisition were filtered to remove common contaminants, IDs originating from reverse decoy sequences and sites originating from the host (human) proteome. MS2/MS3 reporter intensities and the total intensity were then log2 and log10 transformed, respectively. Log2 reporter intensities for each sample were subsequently normalised (centered) by subtracting the median log2 reporter intensity value calculated for all non-phosphorylated peptides detected in the same sample. Data were then filtered by 1 valid value to retain only the quantified phosphosites. Finally, log2 fold changes were calculated relative to a 0s (DMSO) control, separately for the MS2 and MS3 data, to obtain per site response to ionophore treatment. For downstream analysis responses obtained by the MS2 and MS3 based quantification were averaged.

DR thresholds were determined in a timepoint-specific manner by calculating the log2FC MADs scores across each WT timepoint (15s, 30s and 60s), and setting 3x MAD outlier thresholds for each (rounded to the nearest tenth: 15s log2FC<-0.5 for DR^DOWN^ and 15s log2FC>0.5 for DR^UP^; 30s log2FC<-0.6 for DR^DOWN^ and 30s log2FC>0.6 for DR^UP^; 60s log2FC<-0.9 for DR^DOWN^ and 60s log2FC>0.9 for DR^UP^). Phosphorylation sites were considered to be differentially regulated if at any given timepoint their log2FC surpassed these thresholds.

CDPK3 dependency, was determined for each phosphorylation site by calculating the log2 ratios of A23187-treated WT and ΔCDPK3 parasites (ΔCDPK3^A23187^/WT ^A23187^) for each timepoint. The resulting ratios were used to calculate the MAD at each timepoint, and the most stringent score was used to set 3X MAD outlier thresholds (rounded to the nearest tenth: log2FC<-0.6 for CDPK3 dependency in DR^UP^ sites and log2FC>0.6 for CDPK3 dependency in DR^DOWN^ sites). A DR site was considered to be CDPK3-dependent if, at any given timepoint, it simultaneously passed the appropriate DR and CDPK3 dependency thresholds.

#### Clustering

Phosphosite log2FC values from the timecourse experiment were clustered using a Gaussian finite mixture model-based method (Scrucca et al., 2016) log2FC values from both the WT and ΔCDPK3 samples were combined, thus the clustering was performed on six dimensions: WT 15s, 30s and 60s and ΔCDPK3 15s, 30s and 60s. The method was restricted to spherical models with equal or unequal volumes (models “EII” and “VII”) and models with up to 11 clusters were tested. The clustering method was applied separately to the sites designated as up-regulated, down-regulated, and CDPK3- dependent.

#### Gene Ontology Enrichment

Each cluster was tested for an enrichment in Gene Ontology annotations using goatools version 0.8.12 (Klopfenstein et al., 2018). The ontology was downloaded from https://geneontology.org on 2019 April 17. All tests were performed using Fisher’s Exact Test and p-values were adjusted for false discovery rate.

#### Motif Analysis

The sequence surrounding each DR timecourse phosphosite, +/-7 residues, was subjected to a motif analysis using rmotifx 1.0 (motif enrichment; Wagih, Reimand, & Bader, 2015) and WebLogo 3.7.1 (Crooks et al., 2004). The analysis was performed for each cluster as well as for the combined sets of phosphosites designated as up-regulated, down-regulated and CDPK3-dependent.

### Measurement of cyclic nucleotide levels in extracellular parasites

Parasites were seeded onto HFF monolayers in T175 flasks. After 24-30 hours, flasks were washed once with PBS, then scraped and syringe lysed in endo buffer (44.7 mM K2SO4, 10 mM MgSO4, 106 mM sucrose, 5 mM glucose, 20 mM Tris–H2SO4, pH 8.2)/ After counting, the parasites were aliquoted into eppendorfs and treated with 50 µM BIPPO, 8 µM A23187 or the equivalent volume of DMSO for the variable timings while maintained at 37°C. The samples were then lysed by adding two volumes of 0.1 M HCl and left on ice for 10 minutes with intermittent vortexing. The levels of cAMP and cGMP levels in the samples was determined using the enzyme-linked immunosorbent assay (ELISA)-based high-sensitivity direct cAMP and cGMP colorimetric assay kits (Enzo Life Sciences). Samples and standards were acetylated in order to improve sensitivity. All samples and standards were set up in duplicate. Absorbance was measured at 405nm using a FLUOstar Omega plate reader. The detection ranges were 0.078 to 20 pmol/ml and 0.08 to 50 pmol/ml for the cAMP and cGMP assays, respectively.

### DAG & Global Lipid Metabolomics

For the DAG and lipid kinetics experiments, parasites were grown for 3 days in DMEM containing 10% FBS and syringe released by passing through a 23-gauge needle. Lysed parasites were filtered through a 10 μm polycarbonate filter, counted and then pelleted (1,800 rpm, 10 min). After washing pellets with DMEM containing 10 mM HEPEs, parasites were aliquoted to achieve 1x10^8 cells per tube and maintained at room temperature. Parasites were shifted to 37°C and allowed to equilibrate for 60 seconds before addition of pre-warmed DMEM containing either DMSO or 8 μM A23187. Parasites were allowed to incubate for the desired time and quenched immediately on dry ice/ethanol for 5 seconds then left on ice. Cells were pelleted (8,000 rpm, 2 min) and washed with 3x with 1 ml of ice-cold PBS. Cells were pelleted, supernatant aspirated and pellets stored at –80°C until required.

Alternatively for global lipidomics, scraped tachyzoite cultures without stimulus were rapidly quenched in a dry ice ethanol bath and placed on ice. Tachyzoites were then needle passed and filtered on ice and pelleted (1800 rpm, 10 minutes) at 4°C. Cells were washed twice with ice cold 1x PBS and then pellets were stored at -80 until required.

### Lipid analysis

Total lipids and internal standards were extracted using chloroform:methanol, 1:2 (v/v) and chloroform:methanol, 2:1 (v/v) in the presence of 0.1 M HCl with periodic sonication. The organic phase was dried under N2 gas and dissolved in 1-butanol. For DAG, total lipid was then separated by 1D-HPTLC using hexane : diethyl-ether : formic Acid, 40:10:1. For global phospholipid analysis including PA, total lipid was spiked with 1 μg PA(C17:0/C17:0) (Avanti Polar lipids) and then separated by 2D-HPTLC using chloroform/methanol/28% NH4OH, 60:35:8 (v/v) as the 1st dimension solvent system and chloroform/acetone/methanol/acetic acid/water, 50:20:10:13:5 (v/v) as the 2nd dimension solvent system (Amiar et al., 2016). All lipid spots including PA were visualised with primulin and scraped. Lipids and additional standards were then prepared for GC-MS analysis in hexane (Agilent 5977A- 7890B) after derivatisation by methanolysis using 0.5 M HCl in methanol incubated at 85°C for 3.5 hrs. Fatty acid methyl esters were identified by their mass spectrum and retention time compared to authentic standards. Individual lipid classes were normalised according to internal standards.

### Parasite protein extraction, SDS-PAGE, and immunoblotting

Intracellular parasites were scraped and syringe released HFFs by passing through a 23- gauge needle. Extracellular parasites were pelleted (8,000 rpm, 10 min) then lysed in an NP40 buffer (150mM NaCl, 0.5mM EDTA, 1% NP-40, 10mM Tris [pH 7.5]) supplemented with cOmplete EDTA-free protease inhibitor (Roche). Samples were incubated on ice for 10 min, then centrifuged at 12,000xg for 10 min at 4°C and supernatants collected. Following the addition of SDS sample buffer, the samples were electrophoresed on 4- 20% Mini-Protean TGX stain-free precast gels (Bio-Rad) then transferred onto nitrocellulose mem- branes using a semidry Trans-Blot Turbo transfer system (Bio-Rad). The membranes were blocked using 10% skimmed milk in PBS containing 0.1% Tween 20 (PBST) and then incubated with rat anti-HA high affinity (1:1,000; Roche) and rabbit anti-T. gondii CAP (1:2,000; Hunt et al., 2019) for 1 hour, followed by donkey anti-rabbit IRDye 680LT (1:20,000; LI-COR) and goat anti-rat IRDye 800CW (1:20,000; LI-COR) for 1 hour. After several washed with PBS, the remaining bound near-infrared conjugated secondary antibodies were visualised using the Odyssey Infrared Imaging System (LI-COR Biosciences, Nebraska, United States).

### Immunofluorescence microscopy

Parasites were seeded onto HFFs grown in chambered coverslip slides (Ibidi). After 18- 24 hours, the chambers were washed with PBS, then fixed in 3% formaldehyde in PBS for 15 minutes. The cells were then permeabilised using 0.1% Triton X-100 in PBS for 10 minutes, then blocked in 3% BSA in PBS for 1 hour. The samples were then incubated rat anti-HA high affinity (1:1,000; Roche) and rabbit anti-T. gondii CAP (1:2,000; Hunt et al., 2019) for 1 hour followed by goat anti-rabbit Alexa Flour 594 (1:2,000; Life Technologies) and donkey anti-rat Alexa Flour 488 (1:2,000; Life Technologies) secondary antibodies along with 5 μg/ml DAPI for 1 hour. Images were obtained using a Nikon Eclipse Ti-U inverted fluorescent microscope using a 100x objective and processed using ImageJ software.

### PDE Pulldown

HFF monolayers infected with WT or HA-tagged PDE1, PDE2, PDE7 or PDE9 lines were scraped, syringe-released and counted. A total of 1x10^7^ cells per condition were pelleted (8,000 rpm, 10 min) then lysed in 10 μl of NP-40 buffer (150mM NaCl, 0.5mM EDTA, 1% NP-40, 10mM Tris [pH 7.5]) supplemented with cOmplete EDTA-free protease inhibitor (Roche) for 10 minutes on ice. Samples were centrifuged at 12,000 xg for 10 min at 4°C, the supernatants collected and the volume adjusted to 100 μl with IP buffer (150mM NaCl, 0.5mM EDTA, 10mM Tris [pH 7.5] and cOmplete EDTA-free protease inhibitor) to give a final detergent concentration of 0.1%. To pull down HA-tagged PDEs, Pierce anti-HA conjugated magnetic beads (5 μl per condition, Thermo Fisher) were washed 3x with IP buffer to equilibrate the beads, after which the diluted lysate samples was added to the beads and incubated for 2 hours at 4°C on a rotating wheel. After incubation, the supernatant was discarded and the beads washed 3x with IP buffer.

### PDE Assay

The hydrolytic activity of immunoprecipitated PDEs bound to anti-HA magnetic beads was measured using the PDE-Glo assay (Promega), a bioluminsescence-based assay which quantifies the amount of cAMP or cGMP hydrolysed by a given PDE. Briefly, the PDE- bound magnetic beads were resuspended in assay buffer and incubated with either 1 μM cAMP or 10 μM cGMP for 1 hour at room temperature and the reaction terminate by the addition of termination buffer. Detection buffer was added and incubated for 20 minutes, followed by the addition of Kinase-Glo detection solution which was incubated for a further 10 minutes. The supernatants were then transferred to a white 96-well plate and luminescence measured using a FLUOstar Omega plate reader. To test the inhibitory effects of BIPPO on the activity of the PDEs, 25 μM BIPPO or the equivalent volume of DMSO was added to the reaction buffer and left to incubated for 10 minutes before addition of cyclic nucleotide.

### Plaque Assays

Intracellular parasites were treated with 50 nM RAP or the equivalent volume of DMSO for 4 hours, after which the media was replaced, and the parasites left to grow for 3 days to ensure efficient turnover of the PDEs in the RAP-treated samples. DMSO- and RAP-treated parasites were harvested by syringe lysis, counted and 250 parasites seeded on confluent HFFs grown in T25 flasks. Plaques were allowed to form for 5 days, after which cells were fixed in 100% ice cold methanol for 2 minutes and then stained with 0.1% crystal violet for 10 minutes to visualise the plaques. Images of the plaques were acquired with a 4x objective using an Olympus CKX53 microscope fitted with an Olympus DP74 camera. Plaque areas were determined using ImageJ software.

## Acknowledgments

We would like to thank members of the Treeck lab for critical discussions as well as Matthew Child for critical reading of the manuscript. We also thank Michael Howell from the High Throughput Screening Science Technology Platform (HTS-STP), which receives Core Funding at the Francis Crick Institute (FC001999), for performing the imaging for the egress assays. This work was supported by awards to M.T and G.A by the National Institute of Health (NIH-R01AI123457) and to M.T by The Francis Crick Institute (https://www.crick.ac.uk/), which receives its core funding from Cancer Research UK (FC001189; https://www.cancerresearchuk.org), the UK Medical Research Council (FC001189; https://www.mrc.ac.uk/) and the Wellcome Trust (FC001189; https://wellcome.ac.uk/). H.F is supported by Core Funding to the Proteomics Facility at the Francis Crick Institute (Francis Crick Institute (FC001999). B.M.I. acknowledges support from a Wellcome Trust Institutional Strategic Support Award to the University of Exeter (204909/Z/16/Z) and, for the purpose of open access, have applied a CC BY public copyright license to any author’s accepted manuscript version arising from this submission. The work performed by C.Y.B., and N.J.K is supported by Fondation pour la Recherche Médicale (FRM, EQU202103012700), Agence Nationale de la Recherche, France (grant ANR-21-CE44-0010-01, Project ApicoLipidAdapt), Région Auvergne Rhône Alpes (Grant AuRA IRICE GEMELI), Finovi program (Apicolipid project), Laboratoire d’Excellence Parafrap, France (grant ANR-11-LABX-0024), LIA-IRP CNRS Program (Apicolipid project), CEFIPRA-MESRI (Project 6003-1), IDEX Université Grenoble-Alpes. The authors declare that they have no conflict of interest.

## List of supplementary documents associated with the manuscript

**Data S1** – Phosphopeptide quantifications and calculated logFCs from A23187/BIPPO treated WT/ΔCDPK3 parasites. Tabs include data subsets that were subjected to thresholding for differential regulation in A23187 and/or BIPPO treatment conditions. Includes data from TMT sets 1 and 2.

**Data S2** – Phosphopeptide quantifications and calculated logFCs for peptides that were (i) differentially regulated following A23187/BIPPO treatment and (ii) were detected during CDPK3 dependency analysis. Tabs include data subsets that were subjected to thresholding for differential regulation and CDPK3 dependency. Includes data from TMT sets 1 and 2.

**Data S3** – Phosphopeptide quantifications and calculated logFCs from WT/ΔCDPK3 parasites subjected to A23187 treatment timecourse. Tabs include data subsets subjected to thresholding for differential regulation and CDPK3 dependency.

**Table S1** – List of proteins (identified in A23187 timecourse) assigned to clusters identified in the Gaussian mixture-model-based clustering analysis. The clusters are listed across four different tabs based on whether they are differentially up- or down- regulated and whether this shows CDPK3-dependency.

**Table S2** – Gene ontology analysis results of proteins found to be differentially up- regulated in A23187 timecourse.

**Table S3** – Gene ontology analysis results of proteins found to be differentially down- regulated in A23187 timecourse.

**Table S4** – Gene ontology analysis results of proteins found to be differentially up- regulated and CDPK3 dependent in A23187 timecourse.

**Table S5** – Gene ontology analysis results of proteins found to be differentially down- regulated and CDPK3 dependent in A23187 timecourse.

**Table S6** – List of primers used in this study

## Notes

### Competing Interest Statement

The authors have declared no competing interest.

### Summary of Updates

3 minor track changes have been overlooked, and not been accepted in the previous submission. This has now been corrected.

